# A spatial genome aligner for multiplexed DNA-FISH

**DOI:** 10.1101/2022.03.25.485845

**Authors:** Bojing Blair Jia, Adam Jussila, Colin Kern, Quan Zhu, Bing Ren

## Abstract

Multiplexed fluorescence *in situ* hybridization (FISH) has emerged as a powerful approach for analyzing 3D genome organization, but it is eminently challenging to derive chromosomal conformations from noisy fluorescence signals. Tracing chromatin is not straightforward as chromosomes lack conserved shapes for reference checking whether an observed fluorescence signal belongs to a chromatin fiber or not. Here we report a spatial genome aligner that parses true chromatin signal from noise by aligning signals to a DNA polymer model. We demonstrate that this spatial genome aligner can efficiently reconstruct chromosome architectures from DNA-FISH data across multiple scales and determine chromosome ploidies *de novo* in interphase cells. Reprocessing of previous whole-genome chromosome tracing data with this method revealed the spatial aggregation of sister chromatids in S/G2 phase cells in asynchronous mouse embryonic stem cells, and uncovered extranumerary chromosomes that remain tightly paired in post-mitotic neurons of the adult mouse cortex. Our spatial genome aligner may facilitate the adaption of multiplexed DNA-FISH by the community.

## Introduction

Eukaryotic chromosomes undergo dramatic compaction and decompaction in the life cycle of a cell, and the dynamic chromosomal structure plays an integral role in a range of nuclear processes such as DNA replication, recombination, repair, and gene transcription^1–5^. In interphase nuclei, different chromosomes generally occupy separate territories with limited intermingling^6^. Within each chromosomal territory, the chromatin fibers are organized into compartments and domains^7–, 9^, driven in part by the ATP-dependent motor protein complex and loop extruder cohesin^10–14^. The complex chromatin structures enable juxtaposition of remote DNA in space and subsequent transcriptional activation of genes by distal enhancers^5, 15–17^. Disruption of chromatin structures underlies a score of pathologies ranging limb malformations, oncogenesis, and heart disease^18–21^. Delineating how chromatin fibers are folded in the nucleus is therefore of fundamental importance for study of gene regulation and other nuclear processes in health and disease.

Multiplexed DNA Fluorescence In-Situ Hybridization (M-DNA-FISH) has emerged recently as a powerful imaging technique for the study of chromatin structure in eukaryotic cells^22–29^. These technologies utilize serial hybridization of fluorescent probes to tens to thousands of genomic loci in the nucleus to detect the folding patterns of chromosomes. By design, fluorescent probes label discrete genomic loci; the chromatin fiber connecting them is not visualized and must be inferred. The inference of physical connection between two discrete signals is the most salient problem facing chromatin imaging to date. Early efforts to multiplex DNA-FISH often found one, three or four fluorescent signals emanating from one genomic region in a diploid cell line expected to produce two signals^30, 31^. Biological copy number variation, chromosomal intermingling as well as poor probe hybridization have been acknowledged to explain missing signals^26, 27, 30^,; sister chromatids and aneuploidy as well as imaging noise have been acknowledged to explain extra signals^26, 29, 30^. If both noise and biological variation can explain any observed scenario, chromatin fibers cannot be naïvely traced by connecting the first immediate spot. In fact, this uncertainty around imaging has led some to forgo tracing altogether and instead tabulate proximal pairs of imaged loci for bulk analysis^26^. When noise appears indistinguishable from true imaged genomic loci, and biological variation at the single cell level confounds expectation, reconstruction of chromatin fibers remains an intractable computational problem.

In the current benchmark for chromatin tracing, the tracing problem is simplified with assumptions about copy number and emphasize the optical quality of detected signals. Explicitly, an expectation-maximization algorithm (E-M) is first tasked to find *k* chromatin fibers corresponding to an *k*-ploid cell^23, 25^. Repeated *k*-times per cell, candidate fluorescence spots corresponding to a genomic region are scored based on signal intensity, proximity to a moving average of downstream selected spots as well as upstream selected spots, and proximity to a chromosome center (ie. the aggregate of many fluorescent loci) determined by *k*-means clustering^23, 25^. Implicit in this approach are two strong assumptions: that the brightness is a measure of detection confidence and that the copy number of a DNA segment is fixed and known beforehand. However, background fluorescence, non-specific probe binding, and even hot-pixels can frequently emit similarly intense focal signals indistinguishable from the true signal. Additionally, looking for a fixed number of chromosomes may fail to capture true biological copy number variations and aneuploidy. To disambiguate these scenarios, we need a new framework for chromatin tracing that leverages yet unused information, improving on the above heuristics.

Here we present a spatial genome aligner that overcomes the above challenges. We reason that while the shape of chromatin fiber is highly variable, it is subject to spatial constraints dictated by polymer physics^32, 33^. In addition to considering optical quality of signals, our algorithm selects the true fluorescence spots corresponding to a DNA locus from a number of candidates by picking the one that best conforms to a reference polymer model of chromatin. Briefly, these restrictions are the genomic distances between two labeled loci, which should be proportional to its spatial separation. We use a polymer model to estimate an expected spatial distance given a genomic distance and compare the observed spatial distance in imaging to this estimated spatial distance as a test of physical likelihood^32^. We evaluate the accuracy of the spatial genome aligner by comparing pairwise distances discovered by tracing against pairwise contact frequencies discovered by Hi-C. We found that our spatial genome aligner can recapitulate patterns of chromatin organization found in Hi-C at multiple genomic length scales (5 kb, 25 kb, and 1 Mb). Moreover, we found our spatial genome aligner uncovered more chromatin fibers than previously reported in published datasets, and discovered these extra fibers are in fact sister chromatids. We show that each pair of sister chromatids usually reside in a spatially separate chromosome territory, but in ∼2% of replicating cells both pairs of sister chromatids coalesce to interact in one convergent territory. We go on to apply our spatial genome alignment to previous chromatin tracing data generated from mouse cortical excitatory neurons, where we uncover patterns of spatial organization of extranumerary chromosomes inside the nucleus.

## Results

### Spatial genome alignment

Chromosomes are linear, flexible polymers that take on convoluted structures inside the nucleus. One simple but robust model for the spatial configuration of flexible polymers is a Gaussian chain^32^. In this model, the polymer is represented as a chain of successive monomers, linked by bonds of approximately constant length *b*. Each successive monomer is allowed to freely rotate with respect to each other. Transitioning from one monomer to another along the polymer chain is to take one step in a three-dimensional random walk. For any two monomers *i* and *j* on this chain, the probability they are separated by a distance *R_ij_* follows a Gaussian distribution (hence, Gaussian chain)^32^:

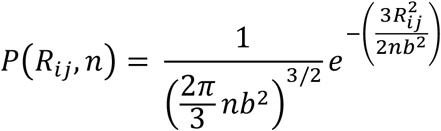

where *n* is the number of bonds, each of length *b*, separating two monomers *i* and *j*.

For a chain with *N* bonds, the likelihood of the entire chain (also known as the conformational distribution function; CDF) is the product of all bond probabilities on the chain^32^:

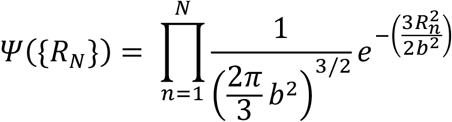

In multiplexed DNA-FISH experiments, we recognize entire chromosomes are labeled at discrete positions, analogous to discrete monomers on a Gaussian chain. Furthermore, these discrete loci are interspaced by regular genomic intervals (eg. 1 Mb), akin to the constant bond length *b* that separate monomers on the model chain. We hypothesized that at large genomic length scales, we can model DNA conformation with a Gaussian chain in which the bond length *b* can be estimated from the genomic distance separating two loci^32, 33^:

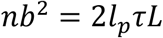

where *l_p_* is the persistence length of DNA, *τ* is genomic-to-spatial distance conversion factor (nanometers per base pair), and *L* is the genomic distance separating two loci, for *l*^*p*^ ≪ *τL*. *τL* together represents the contour length along the DNA polymer separating two loci.

In a setting where multiple signals are detected for *each* of two genomic loci, it is ambiguous which pair of signals lies on the same chromatin fiber. This Gaussian chain model allows us to express the probability two discrete loci imaged are physically connected as a function of both its observed spatial distance and its expected spatial distance. Here, the expected spatial distance is derived from the known genomic distance separating two loci on a reference genome. In taking one step along the chromatin fiber, we can select or omit a fluorescence signal by identifying (if any) a pair of signals whose observed spatial separation is ideally congruent with its expected spatial separation. In tracing the entire chromatin fiber, the most likely polymer among imaged loci is one where the collective segment lengths along the chromatin fiber best aligns with its expected segment lengths. Our optimization objective is therefore to find the sequence of spatially resolved genomic loci that maximizes the likelihood, or CDF, of the polymer traced.

Algorithmically, we first abstract imaged chromatin fluorescent signals as nodes in a directed acyclic graph (DAG). The topological order of nodes is determined by the order of loci on the reference genome (Fig. 1). We connect each node to the adjacent nodes on the linear genome, with each directed edge emulating a polymer segment. For each directed edge, we leverage known genomic distances separating the two imaged loci to estimate an expected spatial distance. We utilize both the expected spatial distance and observed spatial distance between the two imaged loci to calculate a bond probability, assigned as the edge weight. Traversing the graph from beginning to end is to find a potential chromatin fiber. Keeping track and multiplying the edge weights traversed, the score of one path reflects its physical likelihood (CDF).

**Fig 1.**
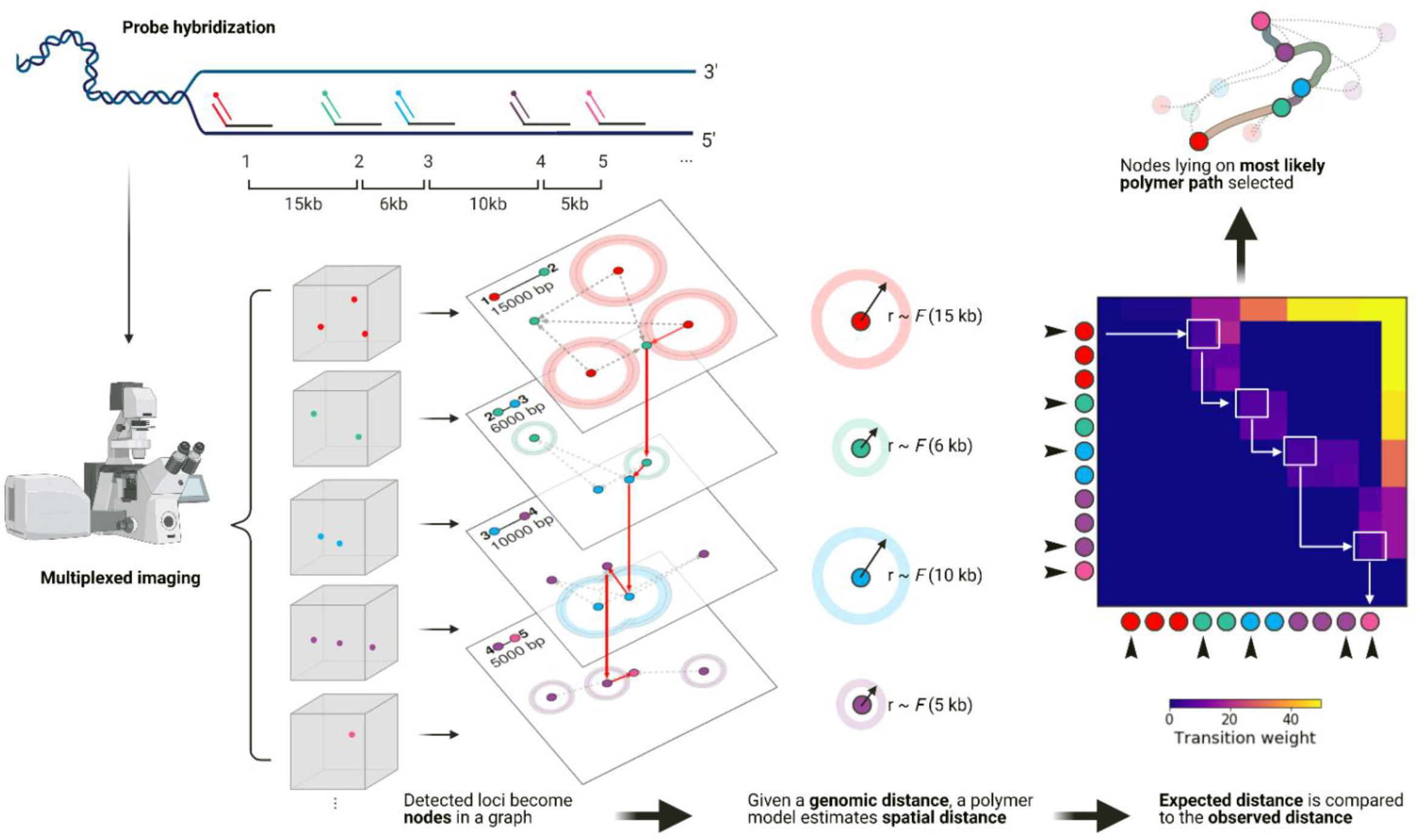
Spatial genome alignment of multiplexed DNA-FISH imaging data against a reference soft-polymer structural model of DNA. Schematic of spatial genome alignment. Spatial coordinates in three dimensions (*x, y, z*) of signal detected from each imaged loci are abstracted as nodes in a graph, ordered by their appearance on the reference genome. Utilizing a freely jointed Gaussian chain model, our aligner estimates an expected spatial distance based on the genomic distance separating two loci. The observed spatial distance is compared to the observed distance, and an edge between two loci are connected weighted proportionally to physical likelihood. We task the aligner to find the shortest path through our adjacency matrix, which returns the sequence of spatial positions whose path length equates to the most likely polymer.

Operationally, we transform the edge probabilities with a negative logarithm function into positive edge weights, such that the additive sum of edge weights reflects the polymer CDF. With this transformation, our optimization objective of maximizing likelihood is reframed as minimizing the sum of negative logarithm transformation of edge probabilities – in other words, we wish to find the shortest path through our graph representation of our polymer. Using dynamic programming, we find the shortest path through the adjacency matrix of our polymer graph not unlike traditional sequence alignment^34, 35^. To account for false positives and false negative imaged spots, all valid paths are explored with the option to “skip” a node permitted by a gap penalty. Since DNA loci from a chromosome must lie on the same chromatin fiber which cannot branch, finding the shortest path is to find the most probable polymer without physical discontinuity discoverable from data^36^.

We first benchmark our spatial genome aligner against the chromatin tracing strategy that connects adjacent genomic loci by converting tabulated distances into an ensemble contact frequency. We analyzed previously published seqFISH+ genome-wide chromatin tracing on mouse embryonic stem cells (mESC), tracing every mouse chromosome at ∼ 1Mb resolution across 1160 single cells^26^. Whereas in published work, detected loci were binned and tabulated to convert distances into an ensemble contact frequency, our spatial genome aligner resolves single-molecule chromatin fibers at single-cell resolution across multiple genome scales. Indeed, our spatial genome aligner traces points whose structures are commensurate with bulk Hi-C (1 Mb: Spearman corr = -0.9 ± 0.04; 25 kb: Spearman corr = -0.85 ± 0.04). We found our spatial genome aligner can resolve large chromatin compartments imaged at 1 Mb intervals as well as finer, single-cell chromatin domains imaged at 25 kb intervals (Fig 2). At 25-kb resolution, local chromatin structure is often nonlinearly organized into topologically association domains (TADs), with sudden shifts in chromatin compaction. Because our polymer model is a freely rotating chain of flexible segments, we found it accommodates such abrupt changes in local topology not easily captured when tabulated in an ensemble fashion. (Fig 2A, B, C).

**Fig 2.**
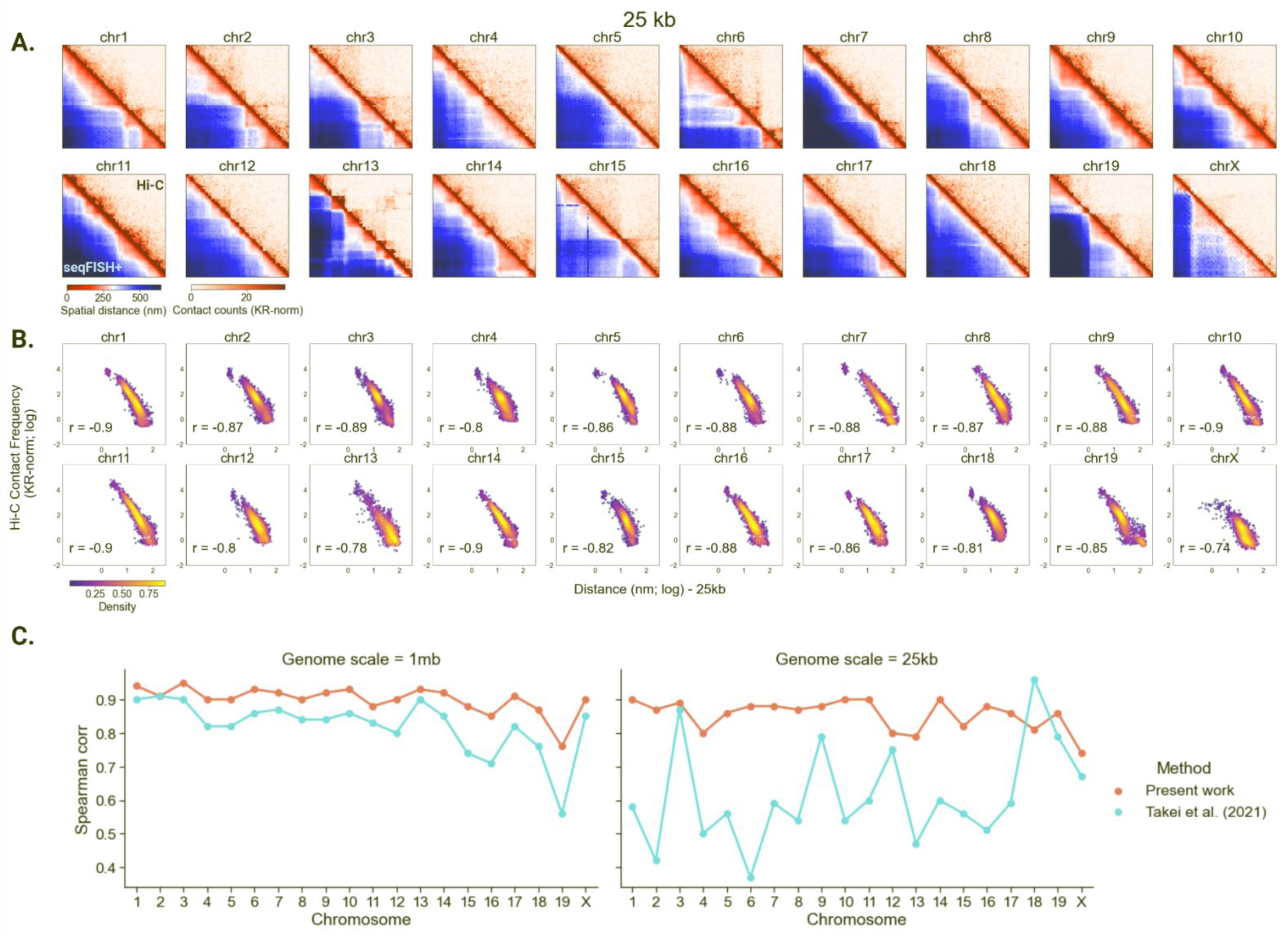
Spatial genome alignment of seqFISH+ chromatin imaging of mESCs at 25 kb and 1 Mb resolution. (A) Heatmaps of seqFISH+ chromatin imaging of mouse embryonic stem cells (mESCs) at 25 kb resolution (bottom left) juxtaposed to contact frequency from bulk proximity ligation assay or Hi-C binned at 25 kb (top right). The distance matrix is an untabulated, median distance matrix of all single-molecule chromatin fibers discovered by spatial genome alignment for every chromosome across 1160 cells. Heatmaps for 1 Mb chromatin tracing imaging data are shown in Supplementary Figure 2. (B) Spearman correlation between pairwise spatial distances (*x*-axis; log-normalized) imaged at 25 kb resolution against Hi-C contact frequency (*y*-axis; log-normalized) binned at 25 kb resolution. (C) Spatial-distance to proximity-ligation correlation comparison across methodologies, at 1 Mb seqFISH+ imaging resolution as well as 25 kb resolution. For 25 kb resolution where DNA domains organize nonlinearly, we note that single-molecule chromatin fiber tracing via spatial genome alignment captures structural variations more faithfully than tabulating all pairwise spatial distances within a specified radius into an ensemble structure.

To evaluate the performance of spatial genome alignment on finer genomic length scales, and on data from other chromatin imaging protocols, we performed spatial genome alignment on multiplexed DNA-FISH data of the Sox2 locus imaged at 5-kb resolution^37^. In our previous work, we adapted a protocol based on sequential DNA-FISH^23^ to label a 210-kb genomic region on mouse chr3, spanning both the Sox2 gene locus in the F123 hybrid mESC line and its super-enhancer 110 kb downstream. By sequentially imaging these loci and tracing the chromatin, we were previously able to visualize promoter-enhancer contacts corralled within a TAD. When we applied our spatial genome aligner to this fine 5-kb resolution chromatin imaging experiment, we found our spatial genome aligner can indeed recapitulate the TAD found at this region (Supplemental Fig 1), faithfully capturing known promoter-enhancer interactions.

We additionally benchmarked our spatial genome aligner with a published chromatin tracing algorithm. Previously, chromatin tracing on multiplexed DNA-FISH emphasized the optical quality of a fluorescence spot, a metric incorporating (a) brightness (b) proximity to a chromosome center and (c) relative agreement to a moving average of preceding and subsequent spots. An expectation-maximization (E-M) procedure then sequentially selected one spot with the highest quality for each chromatin locus, while iteratively updating its quality scores. In contrast, our spatial genome aligner introduces another metric into spot selection – physical constraints dictated by polymer physics – as a decision criterion for selecting spots. Compared to previously published E-M spot selection algorithms (Spearman corr = -0.73), our spatial genome aligner achieves similar accuracy with respect to Hi-C (Spearman corr = -0.76). Notably, our spatial genome aligner performs a global optimization that incorporates the relative positioning of all imaged loci rather than a local moving average, and does so in quasilinear time using dynamic programming. Taken together, we find that the genomic distance separating imaged loci critically helps disambiguate spot selection. Our spatial genome aligner is capable of accurately resolving chromatin fibers at multiple lengths scales, on multiple datasets, and on different multiplexed FISH imaging modalities.

### Polymer Fiber Karyotyping

A nucleus may have multiple copies of a chromosome. Finding all copies has traditionally relied on identifying compact clusters of imaged loci, aggregating by chromatin fiber. A *k*-means approach of clustering assumes the ploidy of a cell is known beforehand^22–27^, and this approach is unable to accommodate copy number variations. For *k*=2 and ploidy *n*=1, *k*-means may inadvertently look for a non-existent second “phantom” chromosome. Conversely, for *k*=2 and ploidy *n*=3, *k*-means would fail to detect an entire chromosome altogether. A ploidy-agnostic approach of clustering, such as DBSCAN, relies on density of detected loci^26, 27^; however, the density neighborhood parameter is difficult to tune. A large density neighborhood may inadvertently aggregate two spatially separable chromosomes as a single dense cluster, misassigning two separate homologs as one. A small density neighborhood may fracture an intact chromosome into separate partitions.

By contrast, our spatial genome aligner provides a density or ploidy independent framework for identifying chromatin fibers. We provide all detected spatial coordinates of a chromosome and a reference genome to our spatial genome aligner, tasking it to extend (if at possible) the most likely path from chromosome start to end. Since the path length (CDF) of a putative polymer reflects the physical likelihood of a polymer, we reasoned that the karyotypes of interphase cells can be obtained simply by counting all physically likely polymer fibers. First, we set a likelihood threshold by scrambling a simulated polymer model of a reference genome such that the observed spatial distances between genes no longer abides by the genomic intervals that separate them (Fig 3B). Next, we iteratively apply spatial genome alignment, extending polymer paths from putative seeds and subtracting nodes visited by the shortest path before searching for next shortest path, until no physically likely polymer path can be discovered (Fig 3A). In this manner, we produce orthogonal sets of coordinates belonging to contiguous chromatin fibers with likelihood scores below our threshold. We call this process polymer fiber karyotyping (PFK).

**Fig 3.**
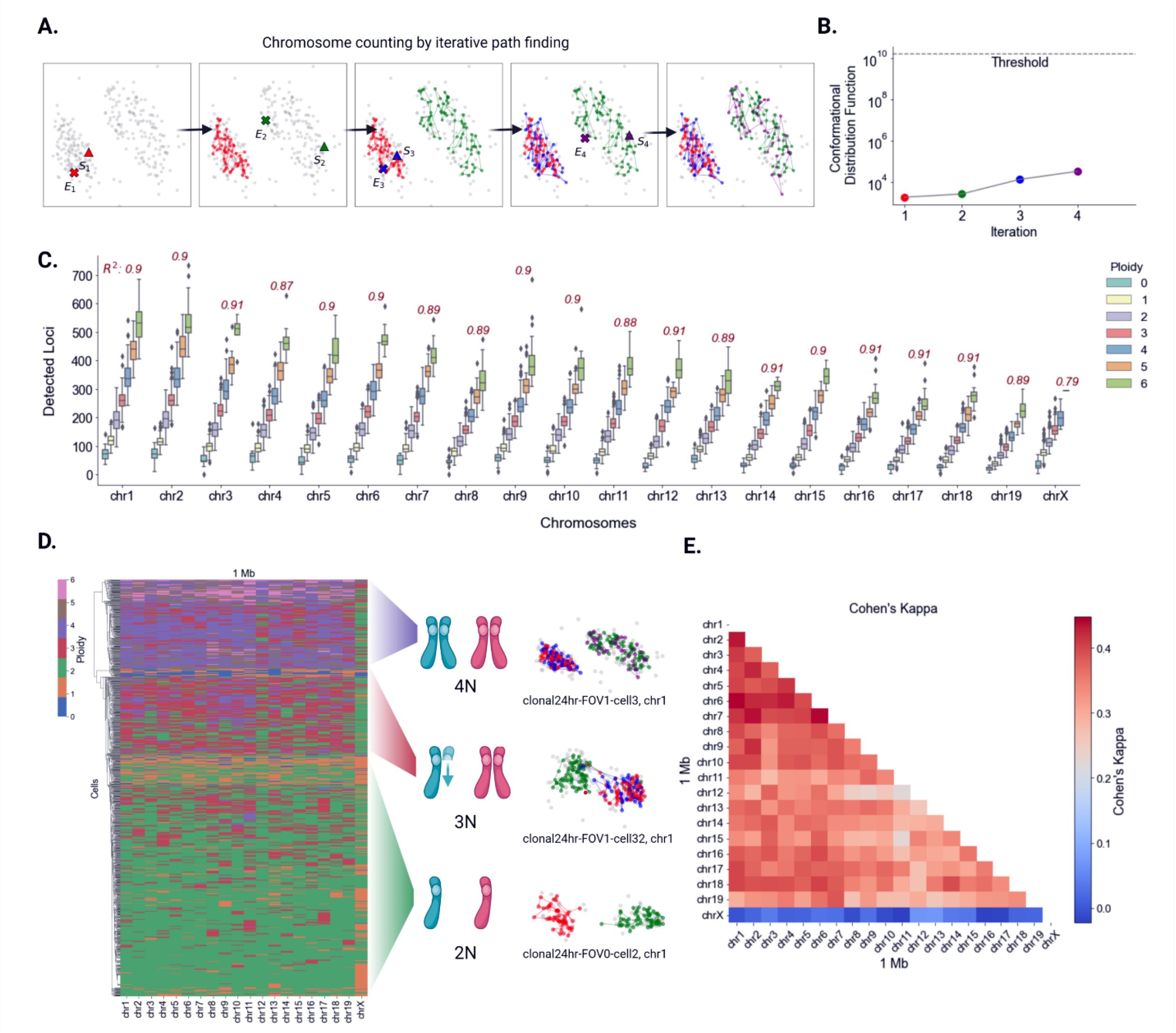
Polymer fiber karyotyping on interphase mES cells. (A) Schematic of polymer fiber karyotyping by iterative path subtraction. For every plausible polymer found, all nodes visited on the polymer path are subtracted before a new graph is constructed for additional rounds of spatial genome alignment. Paths are extended until no physical plausible paths can be discovered. (B) The score of each path is recorded, interpreted as the conformational distribution function of the polymer, and compared to a physically unlikely threshold. Paths are extended for a given chromosome in a cell until either no more paths can be extended or a path extended have a score above threshold. (C) Boxplot of assigned karyotype and total detected loci per chromosome (including spots omitted by spatial genome alignment) of mESC. For every extra chromosome detected by polymer fiber karyotyping, we find a stepwise multiplicative increase in the total detected loci (eg. 1 chr – 100 points, 2 chr – 200 points, …). Pearson correlation coefficient evaluates this trend of detected loci per increase in assigned ploidy. (D) Hierarchical clustering of mESCs by karyotype similarity. Copy number of chromosomes is congruent across the mouse genome in a given cell with the exception of chr X in this male cell line. A dominant faction of cells is 2N in addition to a smaller faction of 3N cell with unsynchronized replication and a smaller faction of 4N cells post-replication. (E) Heatmap of pairwise comparisons of karyotype assigned by different chromosomes, with agreement scored by Cohen’s kappa.

Using chromatin tracing data spanning the mouse genome at ∼ 1 Mb intervals, we performed spatial genome alignment to discover all possible chromatin fibers in the mouse ES cells. Intuitively, a diploid cell should have half as many chromatin fibers as a tetraploid cell; we reasoned this should also reflect in the total number of loci detected in a cell. Comparing the total detected fluorescence signals per chromosome in a cell to its assigned ploidy determined by PFK, we notice a linear relationship. Every incremental increase in ploidy is accompanied by a stepwise, multiplicative increase in the total number of detected loci (Fig 3C). Building on this, we compare the agreement of karyotype assigned by each chromosome. Hierarchical clustering of karyotype assigned by each chromosome across 1160 cells show three distinct clusters of cells whose karyotype are homogenously congruent for all 19 somatic chromosomes. Namely, we see cells proportionally falling into a 6:2:2 distribution of 2N:3N:4N: cells respectively, matching a replicative profile of highly dividing mES cells (Fig 3D). Treating each chromosome as a separate agent for karyotyping, we quantified the karyotype agreement between different chromosomes using Cohen’s kappa test. Pairwise comparisons of each chromosome against another show demonstrably significant agreement (kappa >= 0.3), save chromosome X (Fig 3E). Although the spatial genome aligner had every opportunity to find as many fibers for chromosome X as it did for other somatic chromosomes, it found half as many copies in this male cell line. This gave us additional confidence that our spatial genome aligner produces accurate cell karyotypes without supervision, and that it discriminates karyotype in interphase where even the human eye cannot distinguish true copy number.

### Aggregation of sister chromatids and homologous chromosomes in tetraploid mESC cells

Of the putative 4N cells karyotyped by our spatial genome aligner, we asked if these are polyploid cells with four separable chromosomes before replication, or diploid cells with two pairs of sister chromatids after replication (Fig 4A, B, C). As sister chromatids are shown to be tightly paired in a parallel fashion^38–41^, we reasoned that if two chromatin fibers reside in the same spatial neighborhoods, they are likely sister chromatids of the same homolog. We performed density-based clustering to assign fibers of every ploidy to homologs. Under a set density parameter, the majority of diploid cells had two spatially resolvable fibers singularly residing in different territories (Fig 4B). Notably, under the same density parameter the majority of tetraploid cells also had not four spatially resolvable fibers but rather two clusters of paired fibers that cannot be parsed by eye or by known clustering algorithms (Fig 4B).

**Fig 4.**
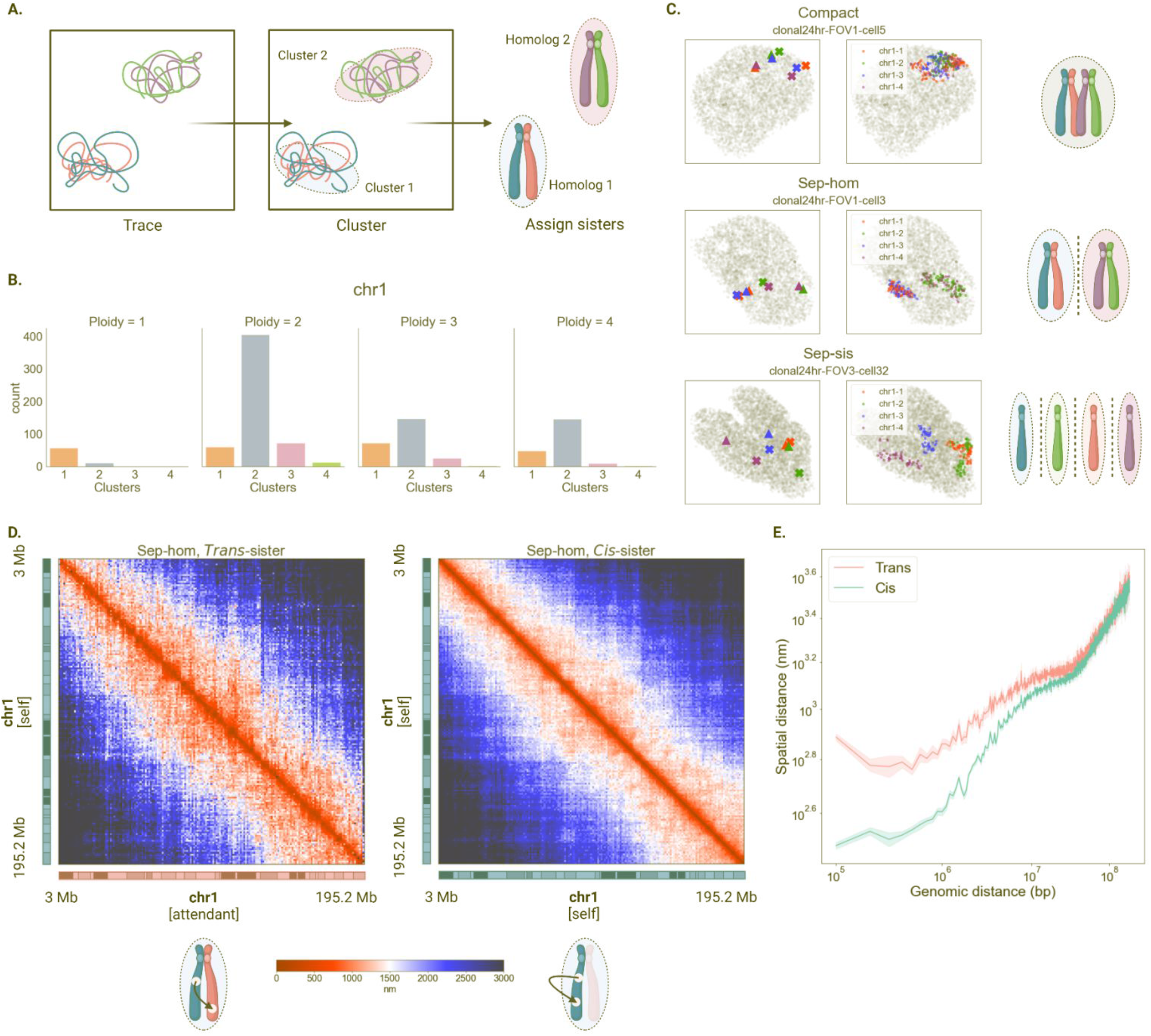
Sister chromatid interactions in mESCs resolved by polymer fiber karyotyping. (A) Schematic of homolog assignment. Spatially separable structures are clustered using DBSCAN and polymer fibers residing in the same spatial neighborhood are classified as sisters of the same homolog. (B) Histogram profile of chr 1 traced at 1 Mb resolution across 1N, 2N, 3N and 4N cells. (C) Representative classes of homologous chromosome organization in 4N mESC cells. A dominant fraction of cells has two spatially separable homologs (sep-hom) residing in chromosome territories, while another fraction of cells has one spatial cluster wherein both pairs of sisters from different homologs interact (compact). A marginal population have three or more spatially separable structures showing separated sister chromatids (sep-sis). (D) Median distance matrices of intra-sister chromatin fiber vs inter-sister chromatin fiber interactions exhibiting tight locus pairing between sister chromatid of the same homolog. (E) Spatial distance separation across genomic length scales between loci on the same sister chromatin fiber (cis) vs separation between loci across different sister chromatin fibers (trans).

To test if these paired fibers are indeed sister chromatids, we examined the *trans-*fiber loci distances relative to the *cis-*fiber loci distances. In agreement with published sister-chromatid sensitive Hi-C on Drosophila and human cell lines^40, 41^, paired fibers in mouse ES cells resolved by our spatial genome aligner are spatially coupled. Explicitly, a given locus of one sister chromatid is followed by the same locus on its attendant sister chromatid, faithfully “shadowing” each other (eg. chr 1, locus 1: µ = 2065.8 nm separation; 90% CI [1801.6, 2330.0]) (Fig 4D). Congruent with previously published work, the spatial distance between *cis-*fiber interactions is closer for smaller genomic distances but which converges with *trans-*fiber interactions above 10 Mb (Fig 4E). Given the parallel nature of pairing and recapitulation of sister-chromatid interactions found by sister chromatid sensitive Hi-C, we presumed the tetraploid cells were in fact replicated diploid cells exhibiting paired sister chromatids.

Because these sister chromatids are tightly coupled, it is plausible loci belonging to one sister are inadvertently selected by its attendant sister and vice versa. To assess this misselection event, we revisited all putative sister chromatid pairs of mESC chr 1. Examining a sliding window of three loci, we allowed sisters to exchange their selected spatial positions for these three loci. Next, we re-evaluated the resultant bond probabilities between three upstream loci to the three downstream exchanged loci (Supplemental Fig 3A). If the new bond probabilities are more probable than the original on both sister chromatid fibers, we call this a misselection event (Supplemental Fig 3B). A sequence of consecutive misselections of any length is called a cross-over event (Supplemental Fig 3D). We found that our spatial genome aligner is susceptible to these errors when tracing paired sister chromatids, and has an average locus misselection rate of 5.06 ± 1.84%. The mean cross-over per fiber is 5.08 ± 3.74 (Supplemental Fig 3D). One source of misselections lies in the sparsity of data (Supplemental Fig 3H). Rarely are two signals simultaneously detected for a locus (7.34%) and present to be selected by both the main sister and its attendant (Supplemental Fig 3H). In fact, in 34.5% of all cases zero signals are detected for a given locus. In the absence of robust signal, our polymer fiber karyotyping routine is greedy. Whichever sister is discovered first is incentivized to select as many detected loci as it sees fits, at the cost of potentially selecting signal that belong to the other sister. A signal cannot be re-selected by subsequent sisters. When only one signal is detected for a given locus, we therefore see a disproportionate imbalance of the single locus signal assigned to the main sister traced first (34%) compared to the signal assigned to the attendant sister traced second (19.6%). Future improvements in experimental detection efficiency, sister-chromatid specific labeling, and algorithmic flexibility permitting revisiting selected signal may decrease this misselection error.

Canonically, homologous chromosomes are divorced from each other in the nucleus and are widely acknowledged to reside in separate territories^6^. Yet, of the 207 cells tetraploid for chr 1 we find two predominant patterns: three-quarters (146/207 cells) bearing two spatially separable clusters presumed to be different homologs (sep-hom), and strangely, a quarter (49/207 cells) with all four fibers are coalesced (compact) (Fig 4C). We note a marginal population of cells (12/207 cells) with three or more separable structures corresponding to separated sister chromatids (sep-sis). In case our clustering density parameter had inadvertently grouped two separable homologs together, we visually inspected each putative compact 4N chr 1. To our surprise, we find a significant proportion of the compact state (31/49 candidates), cumulatively 2.67% of total cell population, has four chromatin fibers spatially intermingling and which cannot be separated by eye.

Why would the newly replicated homolog chromosomes coalesce? We began by parsing which two fibers belong to one homolog within the compact structure. Spatial proximity prohibits clustering from separating homologs and assigning pairs of fibers as sisters. Since true sister pairings should involve two fibers shadowing each other, we reasoned sisters can be assigned by proximity of a given locus on two fibers. There are two natural assignments: grouping the closest pairs by the starting locus (SA; mouse centromere), and grouping the closest pairs by the end locus (EA; mouse telomere) (Fig 5A). We note spatial proximity may cause our spatial genome aligner to inadvertently select spots belonging to other fibers. Therefore, we explored the centromere-centromere distances as well as telomere-telomere distances of the two remaining permutations. Specifically, these permutations correspond to the best possible alternate pairing (alt1) as well as the remaining pairing (alt2), ranked in this order.

**Fig 5.**
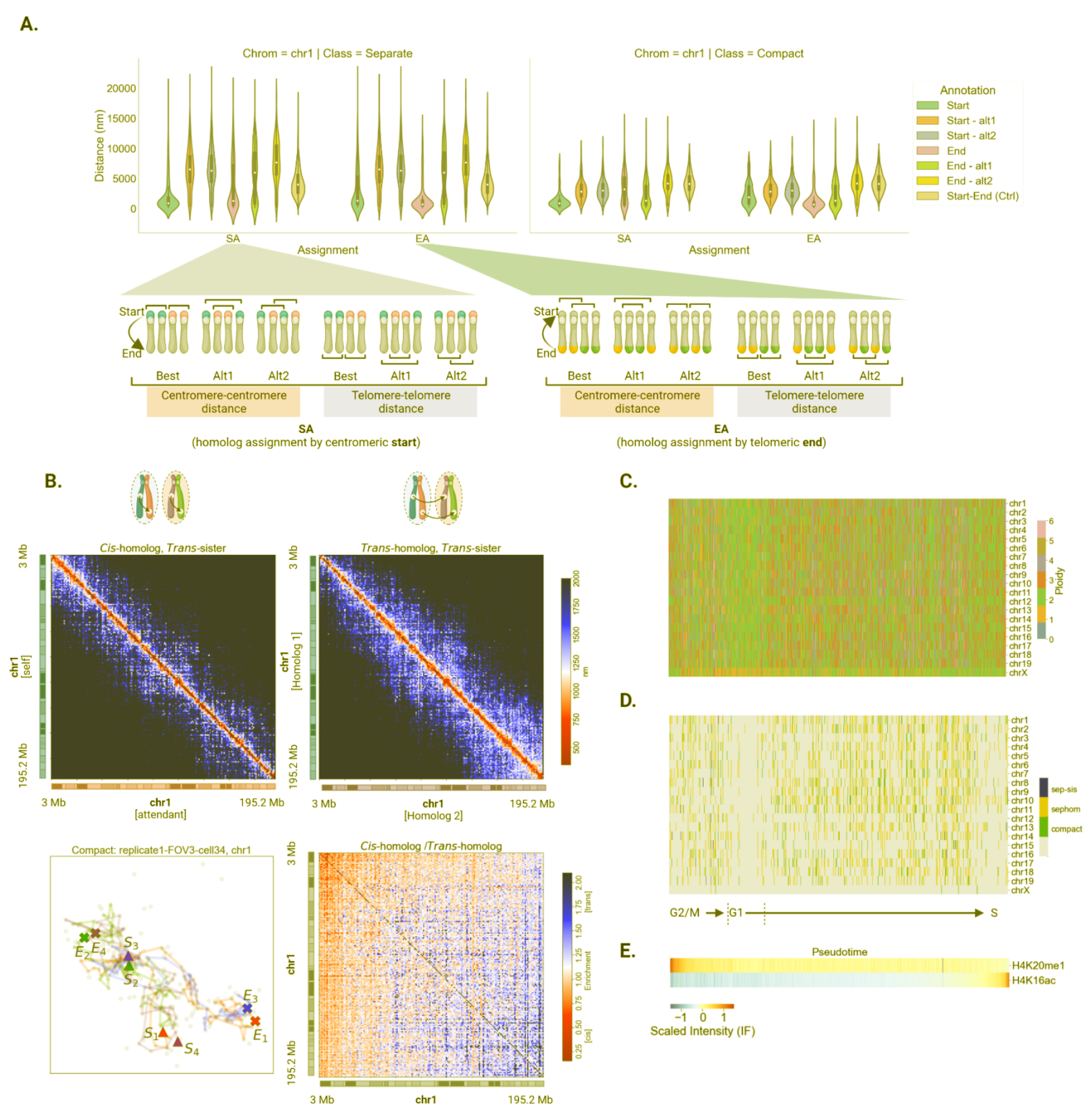
Sister chromatid interactions in compact homolog state. (A) Comparisons of spatial separation between centromeric starts or between telomeric ends of sister chromatids. All 6 permutations, including sisters assigned by closest centromeric starts and its alternate pairings are shown, as well as sisters assigned by closest telomeric ends and its alternate pairings. The pairwise distance between a centromeric start and its telomeric end on the same fiber is presented as a control for unpaired loci. (B) Median distance matrix of *cis-*homolog *trans-*sister chromatids vs *trans-*homolog *trans-*sister chromatids. A representative structure is shown on the bottom left, with a heatmap showing *cis-*homolog to *trans-*homolog distance ratio on the bottom right. (C, D & E) A heatmap of assigned karyotype (C) and tetraploid chromosome organization state (D) ordered along pseudotime determined as a function of cell cycle markers H4K20me1 and H4K16ac (E).

When replicated homologs reside in separate chromosome territories, we find that the telomeres of putative sister chromatids grouped by their centromere (SA) are likely coupled (Fig 4D, left). The mean distance separating telomeres of SA sisters (µ = 3857.5 nm; 90% CI [3451.5, 4263.5]), compared to the distance between its centromere-telomere not known to interact (µ = 4474.9 nm; 90% CI [4295.8, 4654.1]). In contrast, the next best alternative pairing has a telomere separation (µ = 5767.4 nm; 90% CI [5342.2, 6192.6]), larger than two non-interacting loci. In the same manner, the centromeres of putative sisters grouped by their telomere (EA) are also tightly coupled. All other alternate pairings exhibit a spatial separation above that of two non-interacting loci.

When replicated homologs coalesce, we find that putative sisters grouped by their centromere may lose pairing at their telomeres. The mean distance separating telomeres of SA sisters is similar (µ = 3286.5 nm; 90% CI [2835.9, 3737.1]), compared to the distance between two non-interacting loci (µ = 4310.9 nm; 90% CI [4110.2, 4511.7]). Curiously, the next best alternative pairing has a telomere separation (µ = 2544.2 nm; 90% CI [2131.2, 2597.1]) closer than the SA assigned telomere distance. Should this be due to misassignment, then all three pairing scenarios should share a uniformly unpaired distance distribution with mean distances above coupling. Yet, there almost always exist an alternate pairing between putative homologs that theoretically should not interact. The same analysis on EA sisters confers an ambiguous result, likely due to this loss of pairing.

The tendency for *cis-*homolog coupling decreases moving away from the centromere, resulting in a “flare-up” of putative *trans-*homolog interactions near the telomere of chromosomes (Fig 5B). Since seqFISH+ labels discrete genomic loci, we are not able to visualize the contiguous polymer physically linking imaged loci. Additionally, seqFISH+ probes do not discriminate homologs. We are therefore unable to definitively show all spots traced by our aligner lie on the same contiguous fiber, and that our assignments by loci proximity are always sisters of the same homolog. However, our spatial alignment analysis in bulk suggests a possible loss of sister pairing towards the telomeric end and increased *trans-*homologous interaction within this compact 4N state. We ordered the cells along the same pseudotime axis as determined using previously published cell cycle markers (H4K20me1, H4K16ac) (Fig 5C, D, E). We found that this compact 4N state is scattered throughout interphase leading up to S phase. Additionally, this compact 4N state is rarely synchronized across multiple chromosomes, and appears to occur stochastically. This suggests *trans-*homologous interactions between replicated chromosomes is likely uncoordinated and not initiated by a particular cell-cycle checkpoint.

### Chromosomal copy number variations in mouse cortical neurons

Aneuploidy has previously been reported in the brain, by both FISH and single-cell sequencing^44–48^. This copy number variation is thought to underlie a functional diversity adequately supporting neural complexity^45^. We applied spatial genome alignment to whole-genome DNA seqFISH+ imaging of 701 fully segmented mouse neurons^27^, all lying in the center *z*-sections of female mouse cortexes derived from 3 biological replicates. Of these intact nuclei, we honed in on excitatory neurons – the predominant cell type in this dataset (n = 458/701 neurons).

With polymer fiber karyotyping, we also find copy number variations in mouse excitatory neurons. While 58.05 ± 6.38% (mean, standard deviation) of a given chromosome is 2N, we find both putative chromosomal deletions (1N: 13.82 ± 3.03%) and putative chromosomal duplications (3N: 19.68 ± 5.08%) (Fig 6A). Due to data sparsity, many loci are absent on putative extranumerary chromosomes. To ascertain these are indeed three separate chromatin fibers and not two fibers parsed into three, we filtered all putative 3N cells seeking chromosomes that are routed through three separate detections per locus for at least three different loci. We note that each detection of a locus under seqFISH+ imaging is not a one-time detection, but in fact a temporal barcode wherein a locus is detected at least four out of five times, at the correct temporal sequence, while satisfying error correction. Even still, we find that 8.72 ± 4.18% of each chromosome have three fibers (3N) under this more stringent criterion. Of all cell karyotypes, we detect two broad patterns in hierarchical clustering – a group of largely diploid cells with multiple chromosome deletions, and another group of diploid cells with multiple chromosome duplications. We validated our karyotyping results by analyzing haplotype resolved single-cell Dip-C sequencing on mouse cortical neurons classified as excitatory^55^ (Supplemental Fig 4A). By counting total reads, as well as inspecting the relative fold change of reads assigned to the maternal vs. paternal haplotype, the majority of sequenced neurons have balanced haplotyped reads (|log2| fold change < 0.9). Interestingly, 10.82 ± 1.31% of every chromosome have twice (|log2| fold change ≥ 0.9) as many reads of one haplotype as the other (Supplemental Fig 4A, B). Not only may this reflect copy number variations prevalent in cortical neurons, it also suggests the extranumerary chromosome is contributed by one haplotype.

**Fig 6.**
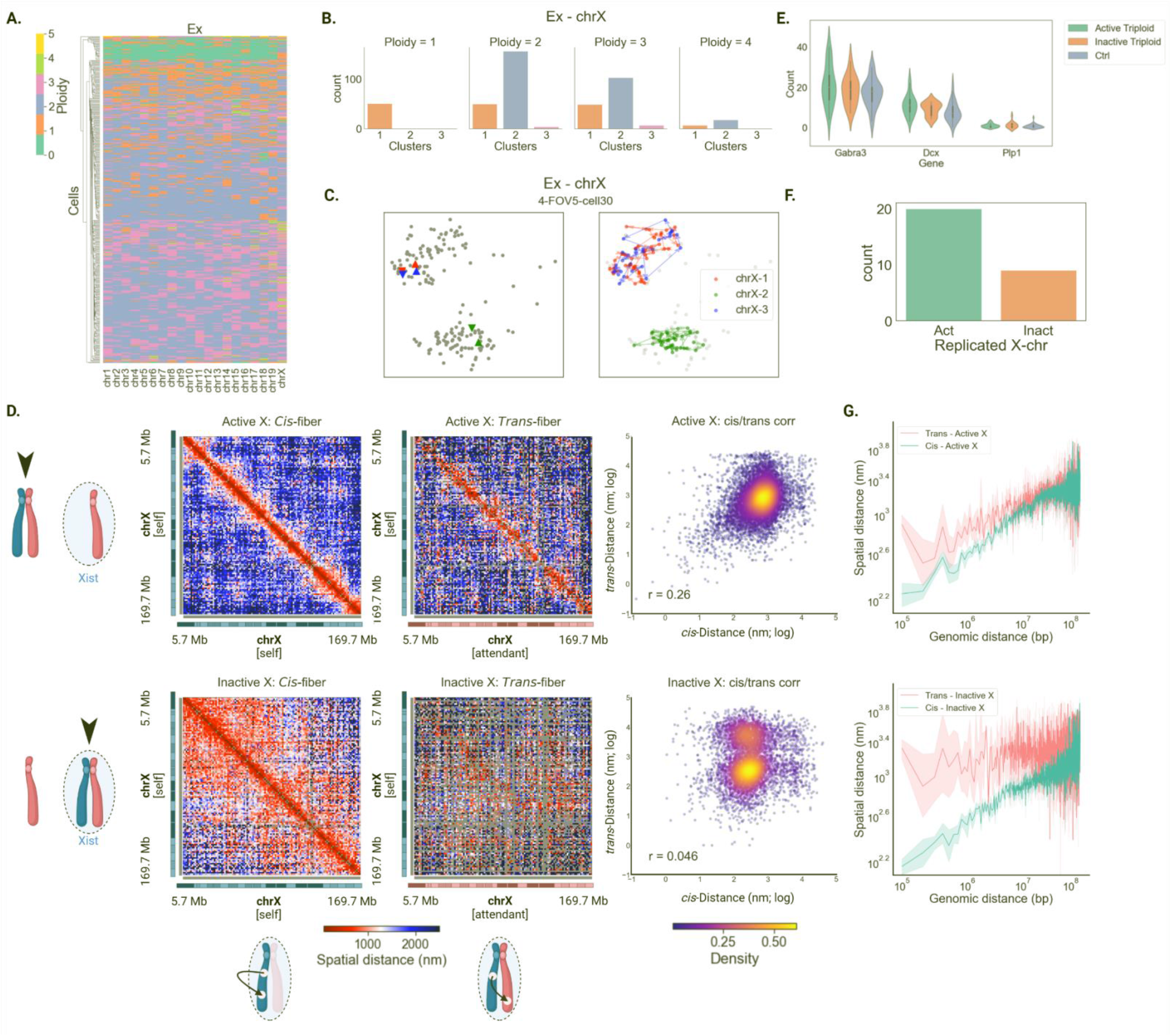
Copy number variations in mouse cortical neurons. (A) Hierarchical clustering of n=458 excitatory cortical neurons from female mice by karyotype similarity. Two dominant factions include a group of cells that are largely diploid with chromosome deletions, and another diploid group with chromosome amplifications. (B) Histogram profile of chr X traced at 1 Mb resolution across excitatory cortical neurons bearing 1, 2, 3, and 4 copies. (C) Representative image of 3N chr X organization in excitatory cortical neurons. Two chromosome territories are preserved despite harboring three chromatin fibers; two fibers reside in one territory while the third is divorced in a separate territory. (D) Median distance matrices comparing *cis-*fiber loci distances to *trans-*fiber loci distances on the active and inactive chr X. Log-normalized distance density heatmap (right) examines the correlation between cis¬-trans distances, on the active and inactive X chromosome. (E) Gene expression counts of three genes lying on chr X, comparing the effect of gene dosage on transcription, sub-classified by chr X activation status. (F) Count plot of excitatory cortical neuron nuclei with three copies of chr X, classified by the activation status of the territory where two of the three chr X fibers colocalize. (G) Spatial distance separation across genomic length scales between loci on the same chromatin fiber (*cis*) vs separation between loci across separate chromatin fibers (*trans*), sub-classified by chr X activation status.

Because single-cell sequencing affords relative copy numbers of reads and fails to capture the nuclear organization of the aneuploid cells, we inspected the spatial organization of these copy number variations. Looking at chr X, where the inactive chromosome is distinguished from the active by RNA imaging of Xist, density-based clustering reveals that excitatory neurons with three chr X have predominantly two chromosome territories (Fig 6B). One chromatin fiber is standalone, while the two remaining fibers are constituents of the same territory (Fig 6C). Of the doubly occupied chromosome territories, two-thirds are devoid of any Xist signal, suggesting a preference for active chr X (Fig 6F). Despite different gene dosage, we detect no significant gene expression relative to diploid cells, irrespective of chr X activation status (Fig 6E). The majority of labeled genes show no significant change relative to elevated gene dosage (Supplemental Fig 5).

Between fibers in the doubly-occupied chromosome territory, the active chr X fibers show some degree of pairing (Spearman r = 0.26) at a locus-to-locus level with its attendant fiber (Fig 6D, G). This manifests as a strong diagonal in the *trans-*fiber pairwise distance matrix, as well as a spatial distance distribution mimicking that of *cis*-fiber pairwise distances. The same cannot be said of the inactive chr X, whose double constituents bear little resemblance (Spearman r = 0.046).

## Discussion

Here we present a spatial genome aligner for multiplexed DNA-FISH data. We showed that this framework resolves chromatin fibers from discretely labeled positions of genomic loci, amid noise and signal dropout. In our spatial genome aligner, each observed locus’ spatial position is checked against a reference model of a polymer chain. This reference model, a Gaussian chain abstracting connections between imaged loci as bond probabilities, dictate that even a highly variable structure as DNA follows predictable patterns of distance separation between loci. Our model accurately captures chromatin compartments and domains on multiple lengths scales and across different chromosomes.

Our algorithm falls into an early lineage of spatial genome aligners, including work by Ross et al. that abstracts connections between loci as polymer segments and whose edge weights are proportional to physical likelihood^33^. Chiefly, whereas a reference polymer structure can reconcile each individual locus’ most likely spatial position using the forwards-backwards algorithm^33, 36^, we demonstrate the utility of dynamic programming to find the most plausible sequence of spatial positions discoverable^36^. In other words, finding the shortest path in our graph representation is to find the most physically-likely polymer without any physical discontinuity. We show finding the most likely contiguous polymer is instrumental to uncovering copy number variations at the single-cell level. Through iterative subtraction of shortest paths finds all valid polymers, we recover sister chromatids otherwise mistakenly grouped as one chromosome fiber. We therefore propose a new form of karyotyping called polymer fiber karyotyping that is density or clustering independent. Ascribing a physical likelihood to polymers allows us to describe in concrete terms detection sensitivity of a copy number of a gene, paving way for the study of copy number variations in interphase for which the expected copy number is unknown. For instance, the study of oncogene amplification in the setting of cancer heterogeneity is currently limited by reliance on compact alignment of probes in metaphase spreads^20^. It is also held back by uncertainty in measurement due to unknown true copy number post-oncogene amplification.

Perhaps not unexpectedly, resolving discrete polymer fibers instead of tabulating chromosome positions uncovers inter-chromosomal interactions. Multiplexed FISH captures in high throughput native chromosome structures directly in intact nucleus among a spectrum of different replicative states. Chromosomes undergo transformative structural change throughout the cell-cycle, disassembling the interphase nucleus to condense into sister chromatids during mitosis – a process that has been intensively studied using proximity ligation sequencing^43^. And yet, to date sister chromatid level interactions have been difficult to resolve from imaging. This is perhaps due to a lack of biochemical labeling that can discriminately label sisters which are identical in sequence. Here, we propose a computational reconstruction relying on statistical mechanics to resolve sister chromatid interactions from fluorescence imaging. We distinguish sister chromatids from a welter of imaged loci difficult to be discerned by the human eye. While the majority of replicated homologs are divorced and reside in separate territories, we did uncover compact territories where all four sister chromatids of a given chromosome spatially aggregate. Whereas in separate territories each sister is shadowed by its attendant sister, in the compact territory sister chromatids can lose sister pairing and might even pair with the other homolog. This structure is evocative of a crossover event, which is thought to occur in mitotic cells at exceedingly rare frequencies (∼1-2%) consistent with our observation^42, 49^. This deserves further investigation to study whether genetic information is indeed being exchanged and may bring significant insight to the regulatory mechanisms of loss of heterozygosity underlying disease. Mechanistically, how a sister chromatid eschews pairing with its own homologous attendant and assiduously choose a sister from the other homolog is a mystery. Additionally, as homologous interactions are highly species specific (ie. Diptera)^38, 39^ we believe this study would benefit from multiplexed DNA-FISH in multiple model systems to study the permeance of this phenomenon.

Multiplexed DNA-FISH affords an intimate look into the elusive inner realities of genomic mosaicism in the brain. We chronicle the intranuclear spatial organization of copy number variations, with single-cell sensitivity and at whole-genome scale, heretofore only reported as frequencies^44–48^. We show that in neurons with three copies of a chromosome, the extranumerary chromosome shares a chromosome territory with another chromatin fiber, preserving two chromosome territories in the nuclei. Within the doubly occupied territory, each chromatin fiber appears to shadow each other, evocative of sister chromatids previously imaged in dividing mESCs. The role and origin of the extranumerary chromosome is unclear. One possibility is that the extranumerary chromosome is the derivative of a non-disjunction event, occurring in a neuroprogenitor during development. Why these chromatin fibers did not dissociate post-division is unclear. Another possibility is that the extranumerary chromosome is a remnant of asynchronous replication, a vestige in a neuroprogenitor that failed to withdraw or complete its replication timing^50^. It has been noted for olfactory receptors, which singularly and stochastically express one receptor gene from an array of over a thousand possibilities, must then also select an allele after a gene selection^51^. The allele selection appears to hinge on asynchronous replication; the expressed allele is faithfully marked by early replication. Others have also noted asynchronous replication as a possible epigenetic mark for monoallelic expression^52–54^. In this spirit and along the emerging field of genome imaging, we believe our spatial genome aligner provides a novel perspective of chromatin imaging analysis with which we hope to unravel foundational biological principles.

## Methods

### Spatial genome alignment algorithm

Conceptually, we abstract each detected fluorescence signal from a 3D image stack as a 4-D node *v = (x, y, z, t)*. Here, *(x, y, z)* correspond to sub-pixel spatial coordinates of a genomic locus detected in imaging, with a resultant positional uncertainty (*σ_x_, σ_y_, σ_z_)* in each spatial axis discovered from 3D Gaussian fitting. The fourth dimension, *t*, corresponds to an order of the gene on the reference genome, ordered from 5’ to 3’ for every chromosome. For notation, we use *v*_*t;i*_ to refer to a node *i* with spatial position (*x_i_, y_i_, z_i_)* corresponding to order *t* on the reference genome; there may be as many as *n_t_* detected nodes for a given gene order *t*.

#### 1. Graph construction

We define a directed acyclic graph *G = (V, E)* as follows:

• *V = {v*_*t;i*_ *}*, *1 ≤ i ≤ n_t_*, *1 ≤ t ≤ T* represents the set of nodes in the graph, for every candidate node *i* of a gene order *t* among *n_t_* candidates, for all *T* genes on the reference genome.

We note that due to signal dropout, there may be genes for which *n_t_ =* 0, in which case no nodes for the given order *t* are populated.

• 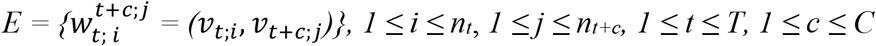represents the set of all edges connecting ordered pairs of nodes, between every candidate node *i* of a gene order *t* among *n_t_* candidates, to every candidate node *j* of a gene order *t+c* among *n_t+c_* candidates, for allowable skips 1 *≤ c ≤ C*, for all *T* genes on the reference genome.

We disallow self-loops by enforcing a lower bound on the skip parameter *c* ≥ 1, such that no edges propagate 3’ → 5’, or more explicitly, no edges from a node of order *t* connect to any node with order less than or equal to *t* +1. This prevents discovery of certain structural variants, such as inversions, translocations, and duplications, but helps restrict solutions to strictly the reference genome. We permit nodes to “look ahead” to downstream genes by skipping up to a permissible upper bound *c* ≤ *C*, scaled later by an affine gap penalty. This accounts for signal dropout resulting in false negative signals, in which case all nodes of a given order *t* maybe false positives and must be skipped.

#### 2. Calculate bond probabilities

We weight the edges using a physical analogy of a polymer model of DNA. Namely, we utilize the freely jointed Gaussian chain model, wherein chemical bonds model connections between two monomers. Here, our discrete spatially resolved genomic locations are analogous to these monomers, connected on the same chromatin fiber. In this model, the spatial distance separating these two locations is modeled after a Gaussian distribution:

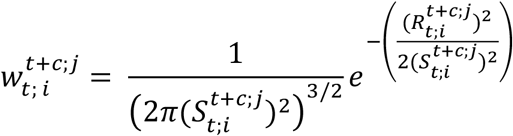

where 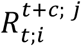 is the distance in nanometers between the *i*th node with gene order *t* to the *j*th node with gene order 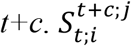 is expanded as:

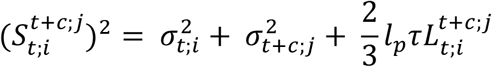

where the positional uncertainties of both the start locus 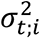 and end locus 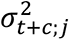 are appended to the second moment 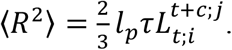.

Here, *l*_*p*_ is the persistence length of DNA in nanometers, *τ* is a scaling factor that converts genomic distance in base pairs to spatial distance in nanometers, and 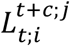 is the genomic distance in base pairs that separate the start locus *v_t;i_*and end locus *v_t+c;j_*. The positional uncertainty is calculated separately for the start and end locus to accommodate chromatic aberrations, in which case loci imaged using different laser channels may have different localization error. 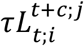 together reflects the contour length of the DNA polymer. Collectively, this allows a comparison of the observed spatial distance 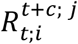 that separate two loci with an estimated spatial distance parametrized as 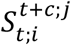 . In this present study, we estimate *τ* by fitting a power-law function through pairwise spatial distance and genomic interval data to estimate a length scale of each base-pair per nanometer.

In this manner, a traversal from chromosome start to end along this graph would accumulate a sequence of bond probabilities whose product reflects the physical likelihood of the discovered polymer:

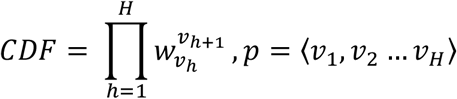

for every node *v* visited on path *p* from source to sink.

Each edge weight 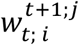 is negative log normalized into positive edge weights whose additive sum is equivalent to the polymer CDF. We do so for several reasons: (a) this controls for numerical underflow in calculating the multiplicative product of small decimals (b) this reframes the optimization objective from maximizing the likelihood function to minimizing the negative log likelihood (c) the CDF is now computed as a sum of edge weights, permitting the use of existing dynamic programming shortest-path algorithms solving for additive edge weights. Below, we write edge weights *w* to represent negative log normalized bond probabilities. To permit nodes to skip potential false positive and “look ahead” to downstream genes, we apply penalize each bond skipped:

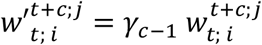

where *γ*_c-1_ is a gap penalty scaled for every skip *c*. Adjacent nodes (*c*=1) are not penalized.

#### 3. Initialize adjacency matrix

We append a single source and a single sink node in our graph to allow gaps in the start and end of polymer alignment. From our graph *G* with total 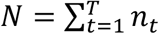 nodes, we construct an (*N, N*) adjacency matrix that is padded by an additional row with index 1 and column with index *N*+2 to an (*N*+2, *N*+2) matrix. We initialize the first row of the adjacency matrix with “pseudo”-bonds that enable up to the first *K* genes to be skipped. These edges linking an imaginary starting position to an observed position are weighted as:

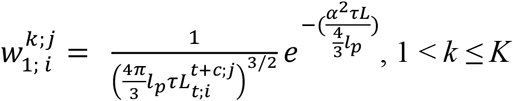

where α scales an imaginary stretched genomic segment with an implicit skip penalty.

We also initialize the last column of the adjacency matrix with “pseudo”-bonds that enable up to the last *K* genes to be skipped. Similarly, the edges linking an observed position to an imaginary ending position is weighted as:

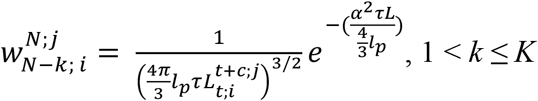

#### 4. Path finding

We find the shortest path from source to sink in our graph using a dynamic programming algorithm. As all edge weights in this directed graph are non-negative, and the sum of traversed edges equate to the CDF of the traversed polymer, we utilize Dijkstra’s shortest-path algorithm as a dynamic programming means for finding the most plausible polymer:

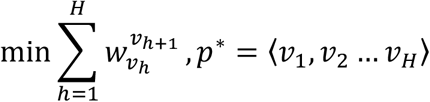

for every node *v* visited on the shortest path *p\**.

We estimate our worst-case time complexity to be:

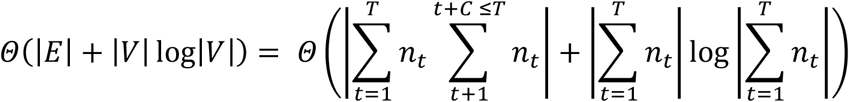

### Polymer fiber karyotyping algorithm

We developed a routine that finds all possible polymers of a given chromosome on a cell-by-cell basis. Chiefly, we accept all polymers below a physical likelihood threshold. This threshold can be derived by scrambling the genomic intervals separating each probed locus, such that the observed genomic distance no longer abides by the expected distances. We then perform an iterative search wherein nodes of each shortest path discovered are subtracted from graph *G* before searching for the next shortest path, until no likely paths below the physical likelihood threshold can be discovered:

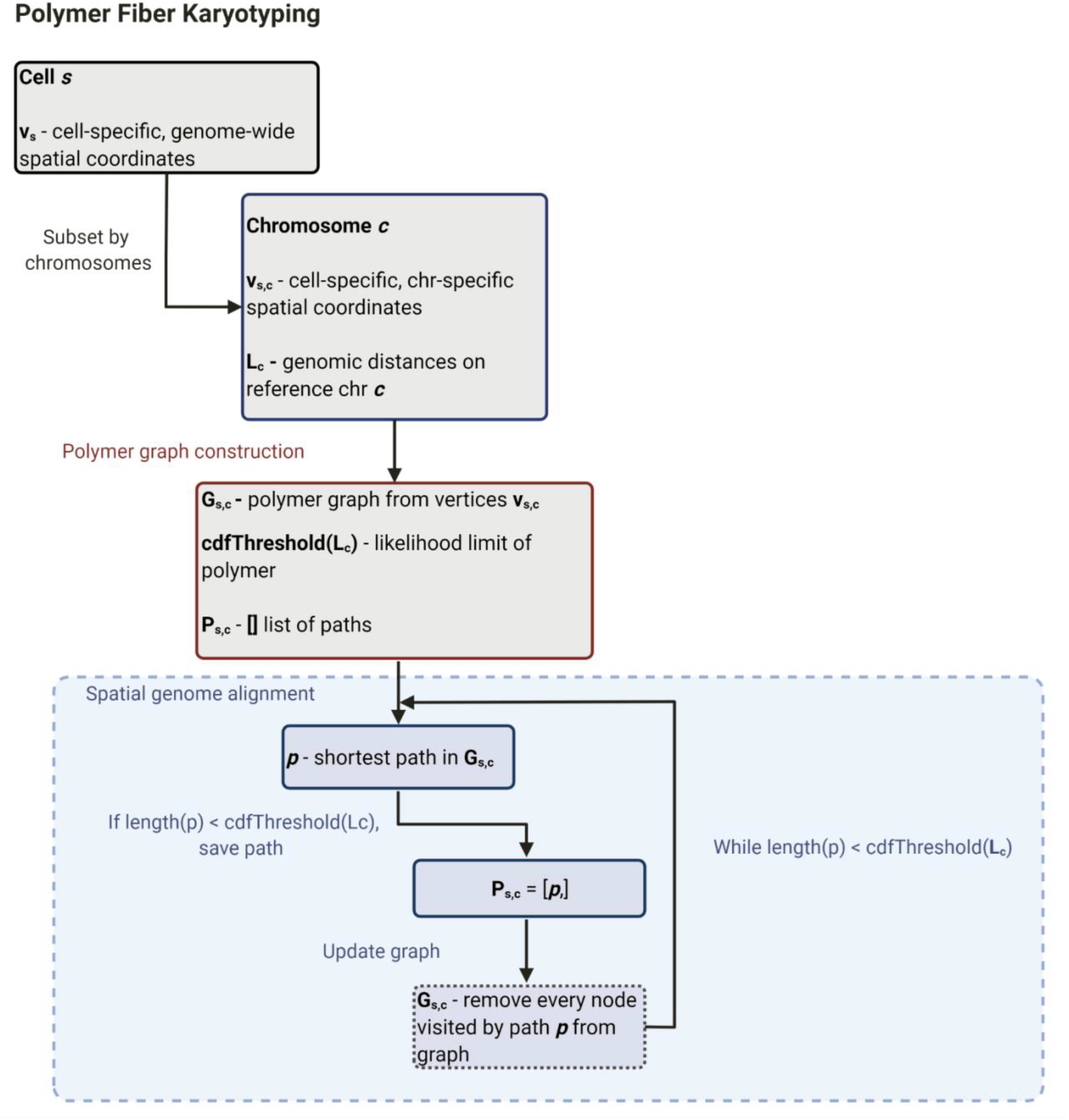

We retrieved spatially resolved fluorescent signal information for mouse genome-wide seqFISH+ probe sets at multiple length scales (1 Mb, 25 kb), and for multiple cell types (mESC: https://zenodo.org/record/3735329; mouse cortical neurons: https://doi.org/10.5281/zenodo.4708112). As previously stated, we fitted a power-law function through pairwise spatial-distances of observed loci, plotted against its genomic-distance separation. From this power-law function, we estimated a parameter corresponding to nanometers per base pair, for every cell-type, for every mouse chromosome. To evaluate the performance of our parameter in its ability to recapitulate true chromatin structure, we performed a hyperparameter search. Using 10% of the total dataset, we performed spatial genome alignment and compared its median distance matrix to Hi-C or cell-type resolved Dip-C. We utilized the parameter that engendered the best fit for our final analysis.

With this spatial distance parameter, for every nucleus and for every chromosome, we iteratively performed spatial genome alignment until no physically plausible fibers could be discovered. Finally, we counted the number of fibers discovered for every chromosome to assign a chromosome copy number, producing a karyotype.

### mESC Hi-C data analysis

To evaluate the spatial genome alignment results of mESCs, we retrieved mESC Hi-C contact matrices from the 4DN Data Portal (experiment set 4DNESU4Y9CBF). Next, we utilized Straw (https://github.com/aidenlab/straw) to extract Knight-Ruiz normalized count matrices for every mouse. Whereas read counts can be evenly binned, the loci imaged by seqFISH+ spanned irregular intervals. To compare Hi-C with seqFISH+ imaging data, where the genomic distances separating each locus are irregular intervals, we performed the following normalization. For seqFISH+ imaging data, we kept the imaged locus ordered 5’ to 3’ closest to each integer 1 Mb or 25 kb bin, dropping all other loci to calculate a final distance matrix. For the corresponding Hi-C matrix, we binned the reads at 1 Mb and 25 kb resolution respectively, dropping the same bins removed from the seqFISH+ distance matrix. To assess spatial genome alignment accuracy, we compared the imaging distance matrix to its corresponding Hi-C matrix using the Spearman correlation coefficient.

### Excitatory mouse cortical neuron Dip-C data analysis

To evaluate the spatial genome alignment results of excitatory mouse cortical neurons, we retrieved cell-type resolved Dip-C contact matrices from NCBI GEO (accession GSE162511)^55^. First, we note a dearth of multi-modal data that ideally allows concomitant cell type classification using one sequencing modality and proximity-ligation analysis on the same cell. In Dip-C, neuronal cell types were resolved by co-projecting *NeuN*+ neurons with bulk Hi-C sequencing of multiple cell-types.

Neurons co-clustering with *Slc17a7+* cells delineating excitatory neurons were also classified as excitatory. It is possible some of these cells classified as excitatory may belong to broader cell types. In contrast, seqFISH+ imaging resolved both RNA and DNA in the nuclei, which allowed excitatory neuron markers (ie. *Slc17a7+*, *Neurod2*) detected in RNA imaging to label cell types. Second, we note a difference in the mouse strains of the two datasets. Dip-C focused on an F1 cross with CAST/EiJ x C57BL/6J background while the seqFISH+ imaging purely focused on C57BL/6J mice.

Amid these caveats, we utilized Straw to extract Knight-Ruiz normalized count matrices for every mouse chromosome. For the seqFISH+ imaging data, we again kept the imaged locus ordered 5’ to 3’ closest to each integer 1 Mb bin, dropping all other loci to calculate a final distance matrix. For the corresponding Dip-C contact matrix, we binned the reads at 1 Mb resolution and dropped the same bins removed from the imaging distance matrix. To assess spatial genome alignment accuracy, we compared the imaging distance matrix to its corresponding Hi-C matrix using the Spearman correlation coefficient.

Using the same Dip-C dataset, we inspected haplotype-resolved reads – namely pre-processed ‘seg’ files – to evaluate read counts assigned to each haplotype. We discarded any ambiguous or multi-contact read-pairs, and counted cells wherein one haplotype has nearly twice as many reads as the other, as a proxy for identifying copy number variations and the haplotype source of those copy number variations at the single neuron level.

### Benchmarking against M-DNA-FISH chromatin imaging protocol and spot selection algorithms

In conjunction, we analyzed M-DNA-FISH imaging of a 210-kb genomic region spanning the Sox2 locus (chr3:34,601,078-34,811,078) based on prior work^23^. We considered all chromosome centers assigned to the 129 allele, which lacks the 7.5 kb tandem CTCF-binding sites (CBSs) inserted on the CAST allele. The previous expectation-maximization routine outlined in (25, 37) generates 10 candidate spots per locus, per putative chromosome in every nucleus, assuming a diploid cell line. Each candidate spot is assigned a score, derived as a combination of (a) fluorescence intensity (b) proximity to chromosome center (c) agreement with moving average of previous loci positions. A candidate spot with a score of -1.5 or more is considered a high-quality spot for the E-M routine, with more negative scores tracking with decreasing quality. For spatial genome alignment, we fed all candidate spots with a much lower quality threshold score of -4, such that for every high-quality spot there is also a low-quality spot. We included this extra noise and ignored the fluorescence intensity information to demonstrate the supreme utility and specificity of genomic distances as a spot selection criterion. We compared the spatial genome alignment results and E-M tracing results against Hi-C, which sequenced the control mouse chr 3 lacking the CBS insertion. We also calculated the Spearman correlation between the discovered pairwise distances and pairwise Hi-C contact frequency for each of the two algorithms.

### Homolog assignment and sister chromatid aggregation analysis

In diploid cells, we utilized density-dependent clustering algorithm DBSCAN (scikit-learn) to separate homologous chromosomes residing in separate chromosome territories.

In tetraploid cells, instead of utilizing density-based clustering to parse and assign homologs, we opted to take a different approach. We first assigned chromatin interaction patterns (ie. separate homologs, compact homologs, separated sisters in tetraploid cells) with DBSCAN to find spatially separable structures and classify tetraploid cells. To assign sister chromatids and in turn homologs, we paired chromatin fibers by the closest starting positions and assigned them as sisters of the same homolog. This allowed us to pair sisters as part of the same homolog in tetraploid cells which had only one spatially dense cluster (ie. compact homologs). In the setting of compact chromosomes where two homologs are spatially not separable, we accounted for alternative pairing scenarios, such as pairing by the telomeric ends. We analyzed the spatial separation of each chromosome starts and ends based on pairing by centromeric starts as well as pairing by telomeric ends, and all possible alternative pairings.

### Sister chromatid misselection assessment

To identify potential loci belonging to one sister chromatid but misselected by the other, we examined all 402 putative sister chromatid pairs for mESC chr 1 (177 loci in length). First, we interpolated spatial coordinates for any loci not detected on a sister chromatid fiber. Next, using a sliding window of three loci beginning at locus 3 and sliding until locus 175, we exchanged spatial positions between the main sister and attendant sister for these three loci. Finally, we re-calculated the bond probabilities of three loci upstream to the three loci exchanged. If the multiplicative product of all recalculated probabilities is more likely than the product of original bond probabilities, we call this a misselection event. We also counted any consecutive sequence of misselections and called this a cross-over event.

### Normalization of chromatin levels for pseudotime analysis

Consistent with previously published work, we constructed a generalized linear model (GLM) for normalizing sequential immunofluorescence data. In this GLM are latent variables controlling for and removing contributions from cell size, total fluorescence intensity over all chromatin marks, as well as batch effect from replicate ID and field of view (FOV). Specifically, we utilized a non-canonical log-link function:

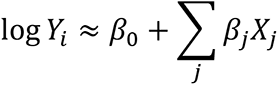

where *Y_i_* is the vector of total fluorescence intensity of chromatin mark *i* across all cells, and *X_j_* are latent variables contributing to bias. Using the Pearson residuals of each fitting, we corrected the readout of two chromatin marks, H4K20me1 and H4K16ac, with which we constructed a principal curve to order cells across the cell cycle or pseudotime. Matching previously published results, H4K20me1 and H4K16ac peaked at opposite ends of the pseudotime axis, ordering cells from G2/M.

### Gene dosage and transcriptional activity analysis

After polymer fiber karyotyping, we stratified cells by the copy numbers discovered for each chromosome and inspected the seqFISH+ RNA imaging corresponding to genes in the seqFISH+ DNA imaging. We performed pairwise t-tests to evaluate statistical significance for the few genes that lie on each mouse chromosome.

### Code Availability

The spatial genome aligner is available at https://github.com/b2jia/jie.

## Acknowledgements

We are grateful for B. Li for bioinformatic support. B.B.J. would like to thank P. Kosuri, V. Bafna, Ross, X. Wen, B. Bintu, Y. Han, Y. Xie, G. Mishne, H. Brandão, E. Mukamel, S. Zu, Y. Zhang, Zhu, Y.E. Li, P.B. Chen, D. Cunningham-Bryant, M. Tastemel and K. Zhang for their invaluable feedback on the spatial genome aligner and polymer fiber karyotyping. The work was supported by NIH Grant 1UM1HG011585 (to B.R.). B.B.J. by was supported, in part, by Ruth Kirschstein Institutional National Research Service Award T32 GM008666.

## Authors’ contributions

This study was conceived by B.R. and B.B.J. B.B.J. designed and implemented the spatial genome alignment and the polymer fiber karyotyping algorithm. B.B.J. analyzed chromatin tracing experiments. B.B.J., A.J., C.K. and Q.Z. benchmarked the spatial genome aligner. B.R. and B.B.J. wrote the manuscript.

## Extended figure legends

**Supplemental Figure 1.**
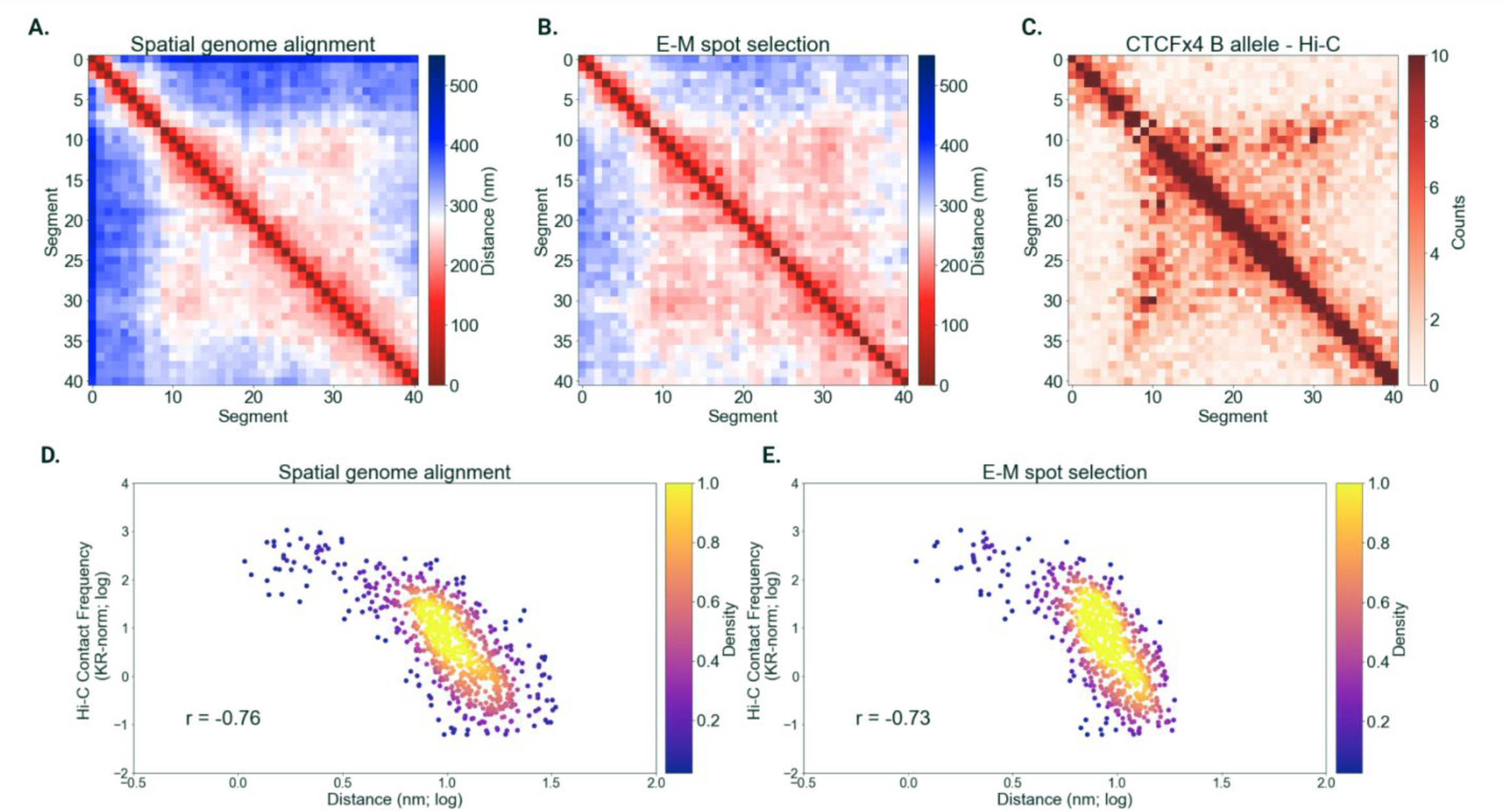
Spatial genome alignment on sequential 5 kb M-DNA-FISH imaging of mESC Sox2 locus. (A) Median distance matrix of spatial genome alignment resolved Sox2 locus (chr3:34601078-34811078), sequentially imaged at a finer 5 kb resolution, using a different imaging protocol based on multiplexed DNA-FISH. (B) Median distance matrix discovered by traditional spot selection using an expectation-maximization routine previously used to analyze M-DNA-FISH datasets. (C) KR-normalized Hi-C contact matrix of the Sox2 locus, juxtaposed as a comparison to median distance matrices discovered by two chromatin tracing algorithms. (D & E) Correlation plot of pairwise genomic distances discovered by spatial genome alignment and an E-M spot selection routine respectively, plotted against KR normalized Hi-C contact frequency.

**Supplemental Figure 2.**
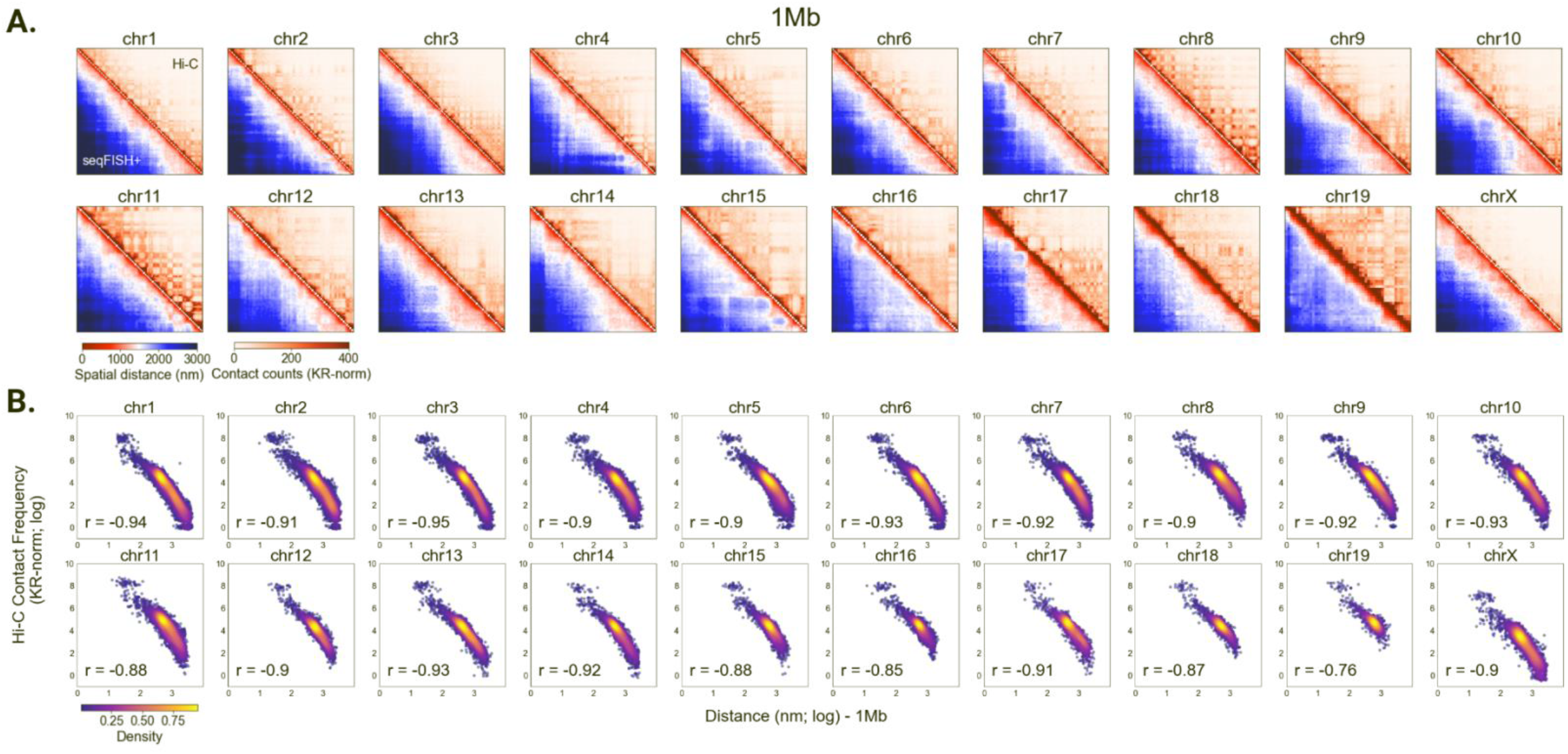
Spatial genome alignment on 1 Mb seqFISH+ chromatin imaging of mESCs. (A) Heatmaps of seqFISH+ chromatin imaging of mouse embryonic stem cells (mESCs) at 1Mb kb resolution (bottom left) juxtaposed to contact frequency from bulk proximity ligation assay or Hi-C binned at 25 kb (top right), continued from Figure 2. (B) Spearman correlation between pairwise spatial distances (x-axis; log-normalized) imaged at 25 kb resolution against Hi-C contact frequency (y-axis; log-normalized) binned at 1 Mb resolution.

**Supplemental Figure 3.**
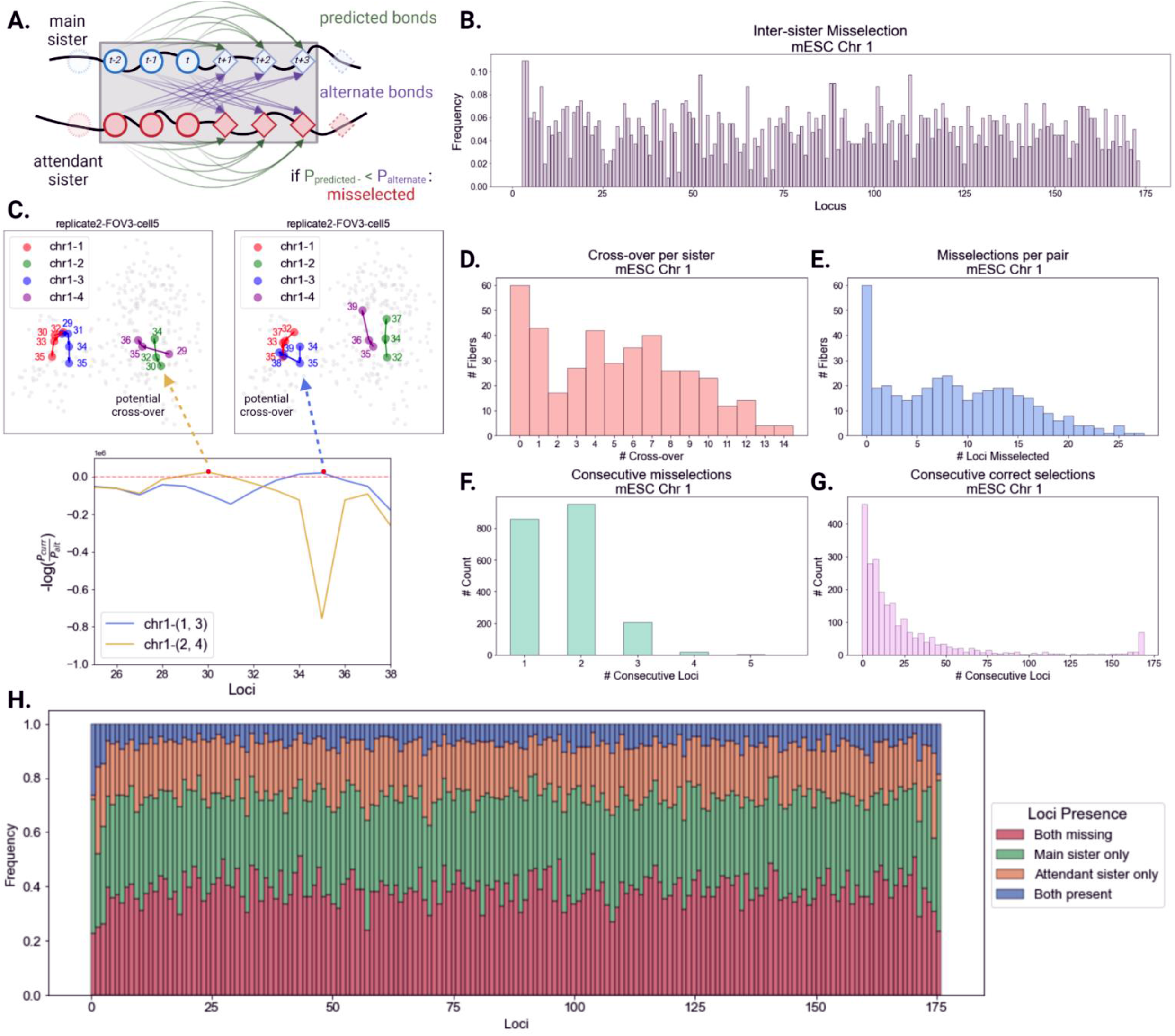
Assessment of misselection between tightly coupled sister chromatids. (A) Schematic of misselection assessment. For every tightly paired sister chromatid, a sliding window of three downstream loci are allowed to exchange with loci on the other sister. Bond probabilities between three upstream loci and three downstream loci are calculated. Original bond probabilities are compared to the cross-over probabilities to evaluate misselection. (B) Bar-plot of misselection frequency per locus on mESC chr 1. (C) Representative image of loci belonging to sister chromatids whose bond probabilities are more probable (-log(P_predicted_/P_alternate_) > 0) if loci are exchanged between sisters. (D) Number of cross-over events, defined as sequences of misselections of any length, per sister chromatid. (E) Number of misselected loci per sister chromatid. (F) Histogram of length of consecutive misselections across tightly paired sister chromatids. (G). Histogram of length of consecutive correct selections across tightly paired sister chromatids. (H) Frequency of loci detection on tightly coupled sister chromatids, stratified by detection on both sister chromatids, detection on main sister, detection on attendant sister, and failure of detection on either.

**Supplemental Figure 4.**
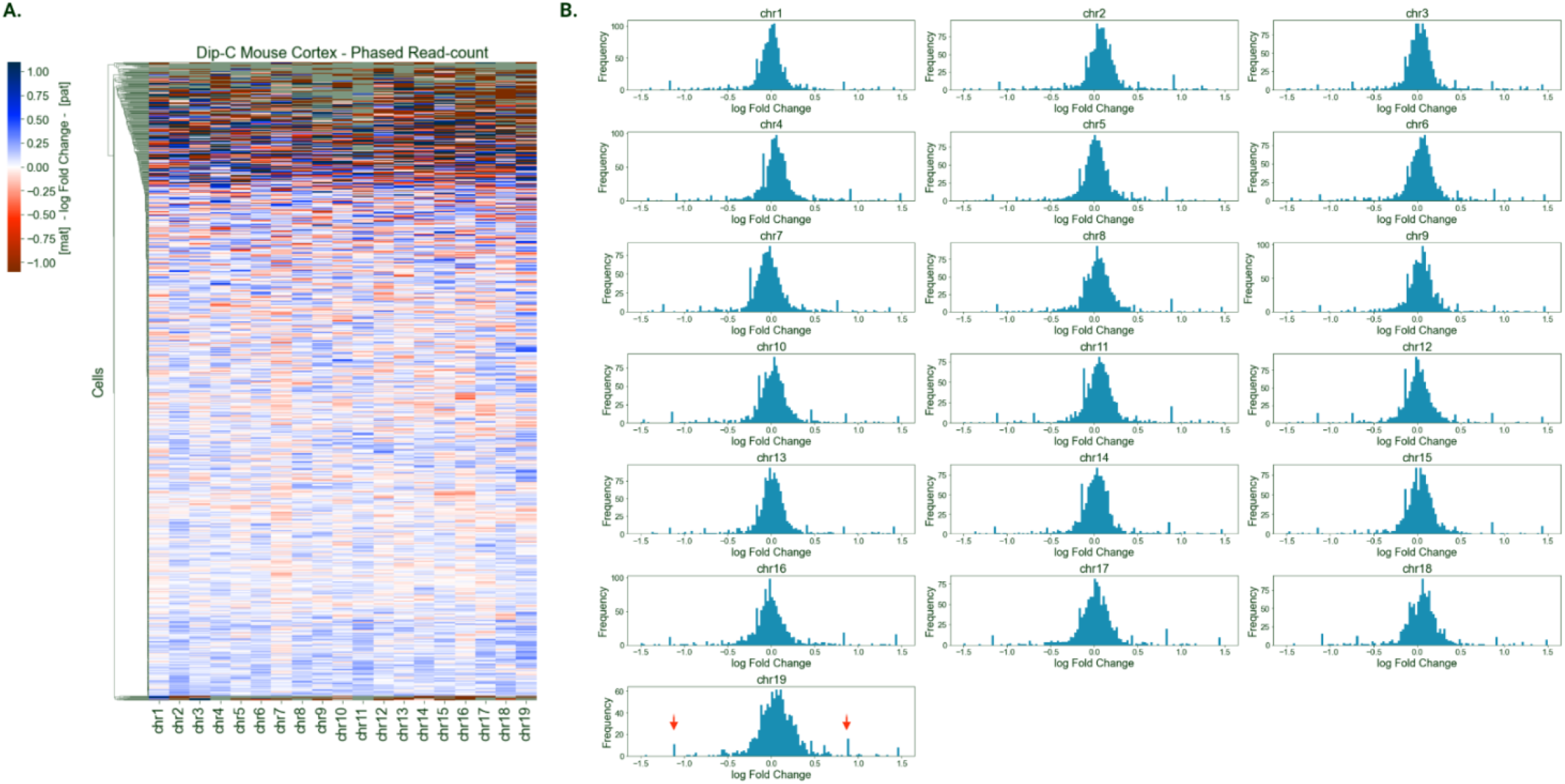
Haplotype-resolved single-cell sequencing of mouse cortical excitatory neurons. (A) Heatmap of log2 fold change of haplotyped Dip-C reads assigned to the paternal vs maternal allele. (B) Histogram of log2 fold change of haplotyped Dip-C reads. Most reads are balanced between the two haplotypes, while a distinct peak around 1 and -1 (red arrows) are present for every chromosome.

**Supplemental Figure 5.**
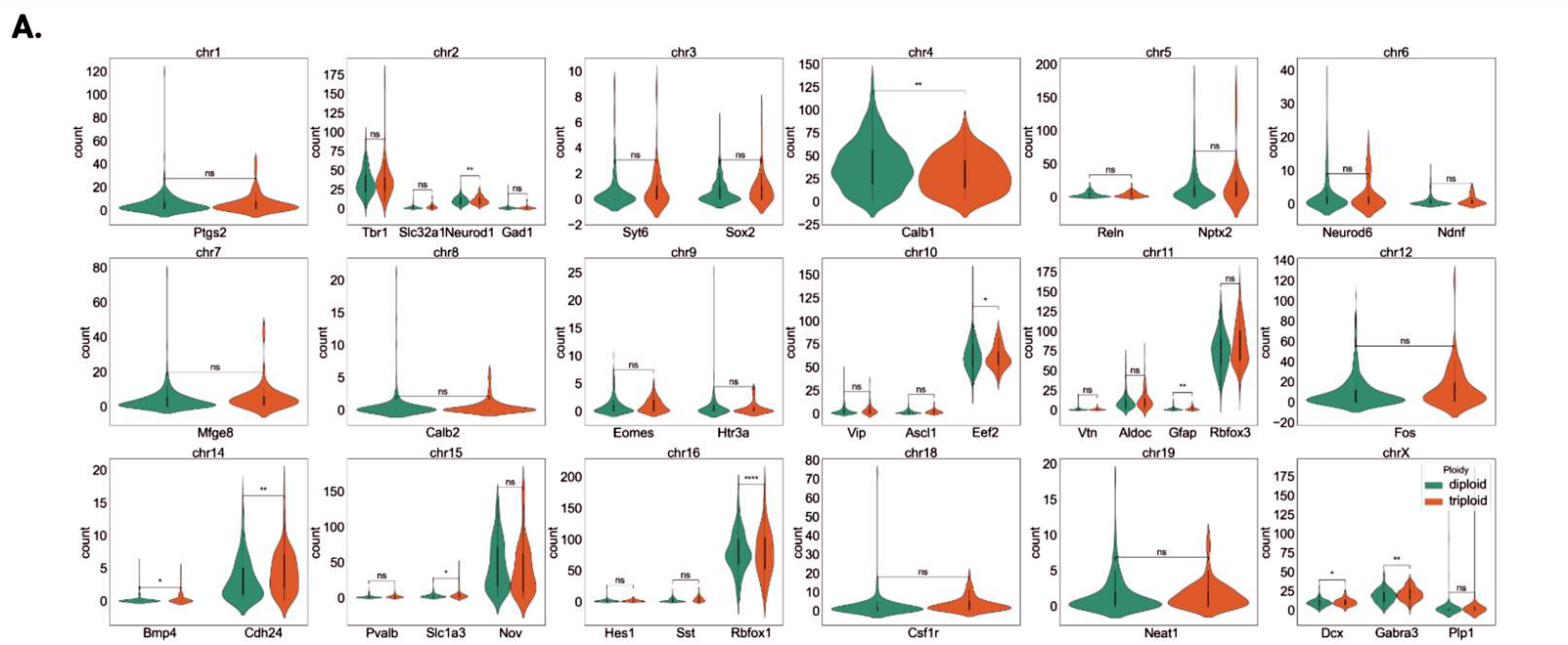
Effect of gene dosage on transcription in excitatory mouse cortical neurons. (A) Violin-plot of RNA transcripts detected by seqFISH+ RNA imaging, corresponding to genes traced by seqFISH+ DNA imaging. RNA counts are stratified by gene dosage, delineated by mouse chromosome. Statistical significance evaluated by pairwise t-tests are labeled above, comparing 2N vs 3N nuclei of a given chromosome.

**Supplemental Figure 6.**
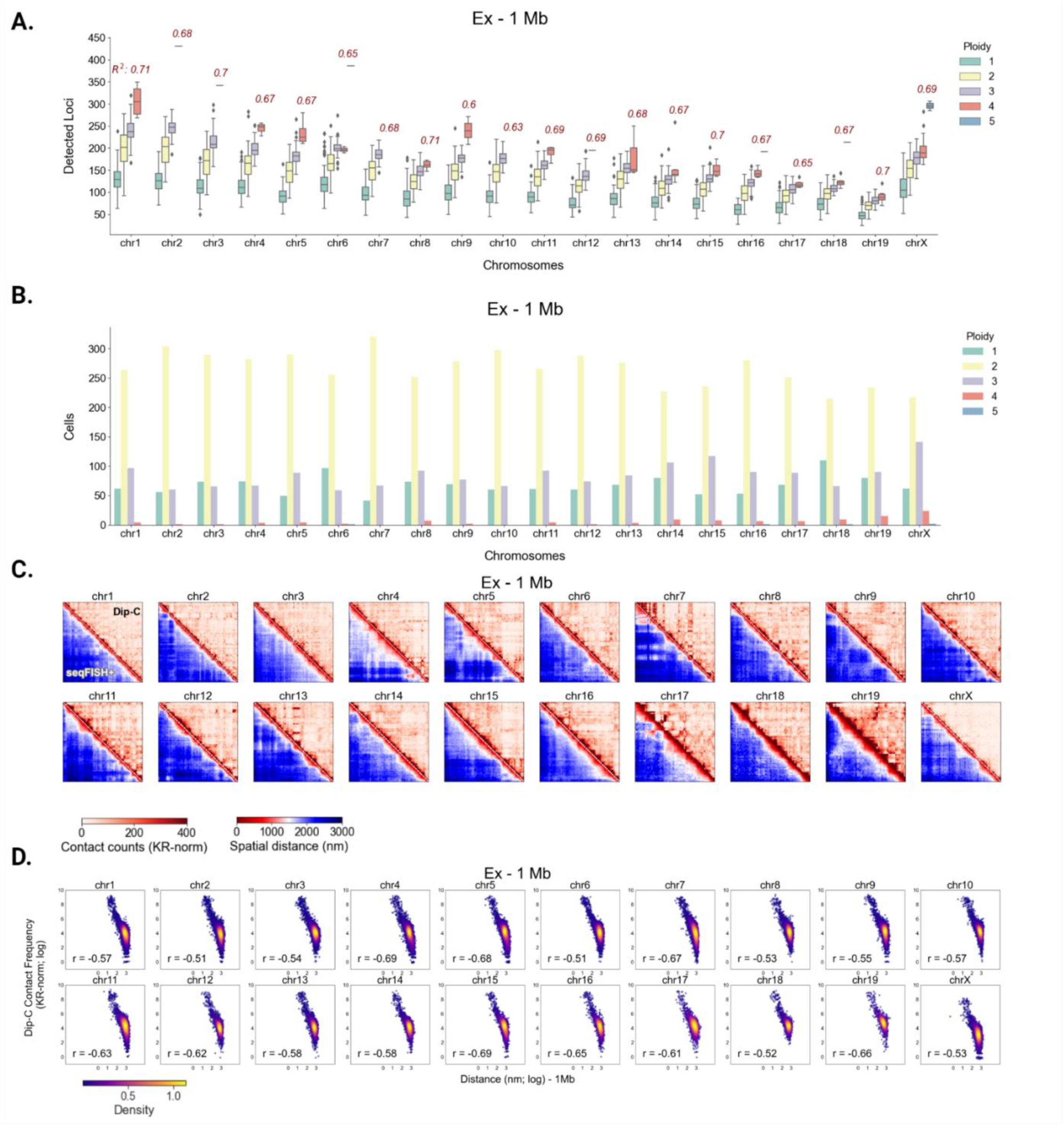
Polymer fiber karyotyping on excitatory mouse cortical neurons. (A) Boxplot of assigned karyotype and total detected loci per chromosome (including spots omitted by spatial genome alignment) of excitatory mouse cortical neurons. For every extra chromosome detected by polymer fiber karyotyping, we find a stepwise multiplicative increase in the total detected loci (eg. 1 chr – 100 points, 2 chr – 200 points, …). Pearson correlation coefficient evaluates this trend of detected loci per increase in assigned ploidy. (B) Histogram of copy numbers discovered by polymer fiber karyotyping on excitatory neurons, delineated by chromosome. (C) Heatmaps of seqFISH+ chromatin imaging of excitatory mouse cortical neurons at 1Mb resolution (bottom left), juxtaposed to contact frequency from single-cell haplotype-resolved proximity ligation assay or Dip-C on excitatory mouse cortical neurons binned at 1 Mb (top right). (D) Spearman correlation between pairwise spatial distances (x-axis; log-normalized) imaged at 1 Mb resolution against Dip-C contact frequency (y-axis; log-normalized) binned at 1 Mb resolution.

